# Differentiated dynamic response in *C. elegans* chemosensory cilia

**DOI:** 10.1101/2022.04.28.489874

**Authors:** Christine W. Bruggeman, Guus H. Haasnoot, Noémie Danné, Jaap van Krugten, Erwin J.G. Peterman

## Abstract

Cilia are membrane-enveloped organelles that protrude from the surface of most eurokaryotic cells and play crucial roles in sensing the external environment. For maintenance and function cilia are dependent on intraflagellar transport (IFT). Here we use a combination of microfluidics and fluorescence microscopy to study the response of phasmid chemosensory neurons, in live *Caenorhabditis elegans*, to chemical stimuli. We found that chemical stimulation resulted in unexpected changes in IFT and ciliary structure. Notably, stimulation with hyperosmotic solutions or chemical repellents resulted in different responses, not only in IFT, ciliary structure and cargo distribution, but also in neuronal activity. The response to chemical repellents results in habituation of the neuronal activity, suggesting that IFT plays a role in regulating the chemosensory response. Our findings show that cilia are able to sense and respond to different external cues in distinct ways, highlighting the flexible nature of cilia as sensing hubs.

## Introduction

Chemosensation is essential for an organism to find a favorable environment. In this fundamental process external chemical cues are detected by receptors on the surface of sensory cells, causing the activation of downstream signaling pathways that eventually evoke a behavioral response. To study chemosensation, we use the soil nematode *Caenorhabditis elegans* as a model organism. This transparent multicellular organism possesses 302 neurons and its entire connectome is known. It is therefore uniquely suitable for studying chemosensing from the levels of individual receptors to sensory neurons and whole-animal behavior.^1,2^ *C. elegans* uses thirty-two ciliated, sensory neurons to detect and respond to the extracellular chemical environment.^3^ Four of these neurons, PHA (left and right) and PHB (left and right), are located in the tail of the animal, with their cilia protruding via the two phasmid channels (left and right) through the animal’s cuticle. These phasmid cilia have been shown to be involved in the detection of repellent chemicals, triggering - together with the amphid cilia located in the head - stereotypical avoidance behavior in so-called drop tests.^4-8^ In a drop test, a solution of a repellent chemical is pipetted on or close to the worm, while its reaction is monitored. In this way, four classical repellents have been identified: sodium dodecyl sulfate (SDS), copper, quinine and glycerol.^3,9^

*C. elegans* neurons function via calcium-based signal amplification,^3,10^ which allows neuronal activity to be monitored using genetically encoded calcium indicators.^11-13^ This approach has been used to study the activity of the amphid neurons.^14,15^ It was found that a variety of chemical stimuli, hyper- and hypo-osmotic shock and nose touch evoke strong activation of these neurons, as was detected by increased intracellular calcium levels, causing depolarization of the cell. In contrast, there are also chemical stimuli that evoke a decrease in intracellular calcium levels, leading to neuronal hyperpolarization. In a recent study using a fluorescent calcium indicator expressed in the phasmid neurons,^16^ it has been shown that PHA and PHB are polymodal sensory neurons, responding to a wide range of chemical stimuli. In that study, worms were glued to a cover glass and exposed to different solutions. In the current study we use a four-flow microfluidics device^17^ to deliver chemical stimuli in a more controlled way, allowing accurately timed exposure to repellents and repetitive switching between repellent and buffer solutions. In combination with fluorescence imaging, this approach enabled us to not only visualize the neuronal responses to the stimuli more accurately and with more detail, but also to monitor the response of components and structure of the phasmid chemosensory cilia at the molecular level.

In *C. elegans*, the sensing machinery, including receptors and other proteins involved in signal transduction, is located in the chemosensory cilia. These are extensions of the dendrites of chemosensory neurons, with a microtubule-based axoneme at the core, enveloped by a membrane.^18^ Cilia do not contain the machinery to synthesize proteins. For their assembly, maintenance and functioning, cilia therefore need a continuous, bidirectional, motor-driven intracellular transport process called intraflagellar transport (IFT).^18-20^ In IFT, trains consisting of multiple IFT-A and IFT-B protein subcomplexes move along the central axoneme to deliver cargo, such as tubulin and receptors for chemosensing, to the tip of the axoneme and back to the base.^21^ Anterograde transport, from ciliary base to tip, is driven by motor proteins of the kinesin-2 family: in the proximal segment of the cilium heterotrimeric kinesin-II is the main motor protein, further along the cilium homodimeric OSM-3 gradually takes over transport.^22^ At the tip of the cilium, IFT trains disassemble and reorganize into retrograde trains, that travel back to the base of the cilium, driven by IFT-dynein.^18,23-25^ So far, it is not known whether and how IFT and ciliary structure are affected by exposure of the chemosensory cilia to chemical stimuli.

Our approach, combining microfluidics for stimulus delivery^17^ with fluorescence microscopy, enables us to study the multiscale process of chemosensation in live *C. elegans*: from the response of the ciliary IFT machinery to neuronal activity and subsequent behavioral changes in response to various repellents. We show that chemosensing not only results in a neuronal and behavioral response, but also in striking changes to IFT and ciliary structure. Our findings highlight that the cilium is a highly flexible organelle that is able to sense and repond to different stimuli in distinct ways.

## Results

### Phasmid chemosensory neurons show distinct activity profiles in response to various repellents

In order to study neuronal activity of the phasmid chemosensory neurons PHA/PHB in response to repellent chemicals, we made use of a worm strain expressing pan-neuronal GCaMP6s,^11^ allowing visualization of calcium spiking, and thus neuronal signaling, in all neurons. We used a microfluidics chip^17^ with a four-flow system for stimulus delivery to stimulate the worm in a spatiotemporally controlled way. By putting the worm in the chip’s worm channel tail-first,^26^ we could specifically deliver stimuli to the tail, where the phasmid chemosensory neurons are located (Figure 1A). Stimulation resulted in a clear increase in GCaMP6s fluorescence intensity in the dendrites and the somas of the PHA/PHB neurons (Figure 1B). Control experiments with buffer in all flow channels showed that changes in pressure did not evoke calcium spiking, demonstrating that the observed fluorescence increase upon stimulus delivery is indeed caused by the chemical nature of the solution.

**Figure 1:**
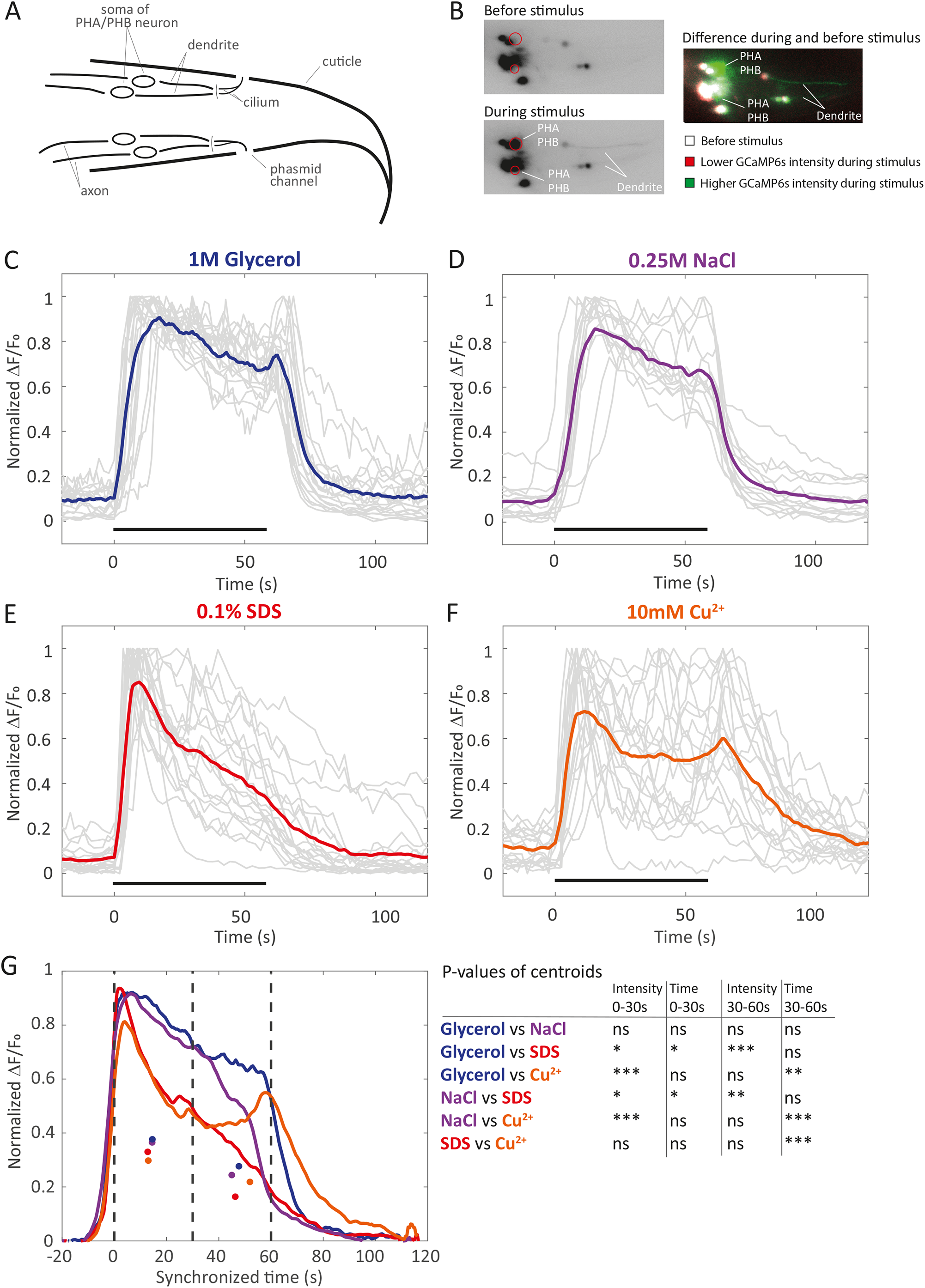
Polymodal signaling of PHA/PHB neurons in response to various stimuli. See also Movie S1, S2; Figure S3. A Schematic overview of *C. elegans* tail, showing the four phasmid chemosensory neurons and their cilia, which are exposed to the environment via the phasmid channels. B Inverted fluorescence microscopy images of the tail of *C. elegans* expressing pan-neuronal GCaMP6s, after exposure to M13 buffer (upper left image) and glycerol (lower left image). Right: difference image of lower left and upper left image, indicating the changes in GCaMP6s intensity upon stimulus exposure. Situation before stimulus is depicted in white; GCaMP6s signals that are lower during stimulus are depicted in red; signals that are higher are depicted in green. Note that positive and negative intensity scaling have been optimized independently for maximimum contrast; false color in right images should only be interpreted qualitatively and not quantitatively. C-F Normalized GCaMP6s fluorescence intensity profiles in the PHA/PHB somas in response to 1M glycerol, n=19-11 (C); 0.25M NaCl, n=14-9 (D); 0.1% SDS, n=21-10 (E); and 10mM Cu^2+^, n=20-13 (F) in M13. Light grey lines show individual measurements, thick colored lines averages. Black horizontal bars indicate onset and duration of the stimulus. n=x-y with x the number of somas and y the number of animals (see Methods). G Averages of normalized GCaMP6s fluorescence intensity profiles in response to 1M glycerol (blue), 0.25M NaCl (purple), 0.1% SDS (red) and 10mM Cu^2+^ (orange) (as shown in C-F) as a function of time (synchronized). Vertical dashed lines indicate the two time intervals that were used to calculate the centroids, which are indicated with a dot in the corresponding color. To compare the traces in response to the four different stimuli, we performed an ANOVA test with multiple comparisons for the centroids (see also Figure S3D). Statistical significance is indicated in the table, ns = not significant; * p<0.05; ** p<0.01; *** p<0.001.

By quantifying the GCaMP6s fluorescence intensity in the PHA/PHB somas over time, we analyzed the neuronal activity in response to aversive solutions (Movie S1A,B). Immediately upon stimulation with glycerol, a hyperosmotic solution that acts as repellent,^7^ GCaMP6s fluorescence intensity increased sharply, remaining high for the entire duration of the stimulus (Figure 1C). As soon as the stimulus was removed, GCaMP6s intensity levels returned to baseline level. Stimulation with a lower concentration of glycerol (0.5M) resulted in similar calcium traces, with high intracellular calcium levels for the entire duration of the stimulus (Figure S3A). Stimulation with high concentrations of NaCl, with an equal osmolality as the 0.5M glycerol stimulus, showed similar GCaMP6s intensity time traces (Figure 1D, Movie S1B). When we increased the duration of the hyperosmotic stimulus, the GCaMP6s fluorescence intensity, showed a fast increase, followed by a gradual decrease, plateauing eventually at ∼40% of the initial level and staying constant until the stimulus was removed. We note that photobleaching was negligible at the imaging condition used (Figure S3B), indicating that the longer-term decrease of the fluorescence intensity is most likely due to a partial decrease of the intracellular calcium concentration during the stimulus. These results show that exposure to hyperosmotic stimuli results in increased intracellular calcium levels for as long as the stimuli last.

In contrast, stimulation with SDS, a detergent that also acts as repellent^7^ (Movie S2), showed different GCaMP6s fluorescence intensity time traces (Figure 1E). After an initial sharp increase in fluorescence intensity, the signal decayed quickly back to baseline level and remained low during the rest of the stimulus exposure. Similar time traces were observed in response to a solution of CuSO_4_(Figure 1F, Movie S1A): immediately upon stimulation with Cu^2+^, GCaMP6s intensity increased sharply and returned to baseline level after the initial peak. Notably, in many measurements a second and sometimes even third peak was observed. When we increased the duration of stimulus exposure, we observed repeated peaks in GCaMP6s intensity in roughly 50% of the cases (Figure S3C). It could be the case that the PHA and PHB neurons show different activity profiles, but it was not possible to distinguish between the two, since we used a worm strain in which all neurons express GCaMP6s.

To provide statistical evidence of similarities and differences in the GCaMP6s intensity profiles in response to the four stimuli, we synchronized the intensity time traces in such a way that the initial increase in fluorescence intensity at the start of the stimulus was set to t=0. We split the stimulus exposure time in two intervals of 30 seconds each and calculated the centers of mass of both halfs of the curves (Figure 1G; Figure S3D). This allowed statistical comparison of the calcium traces obtained in response to the four stimuli. In the first 30 seconds of stimulus exposure, the GCaMP6s intensity levels fall in two statistically distinct groups: on the one hand the responses to glycerol and NaCl are similar, on the other hand the responses to SDS and Cu^2+^. At their centroids, SDS and Cu^2+^ show lower GCaMP6s intensity levels than glycerol and NaCl, which is consistent with the observation that the decrease in GCaMP6s intensity after the initial peak that we observed for SDS and Cu^2+^ is not present for glycerol and NaCl. In the second half of the stimulus exposure (from 30 to 60 seconds), the GCaMP6s intensity levels at the centroids are significantly different for SDS on the one hand and glycerol or NaCl on the other. The GCaMP6s intensity for Cu^2+^ in te second interval was not statistically different from glycerol and NaCl, most likely due to repeated, additional peaks in GCaMP6s intensity we particularly observed in this condition. These peaks explain why the centroid for Cu^2+^ is shifted to later time compared to the other three stimuli. An ANOVA test with multiple comparisons for the centroids showed that the response to Cu^2+^ is more similar to SDS than to glycerol and NaCl (Figure S3D).

Taken together, stimulation with a hyperosmotic solution, such as glycerol or high concentrations of NaCl, led to prolonged increased intracellular calcium levels, for the entire duration of the stimulus. Upon stimulation with SDS or Cu^2+^ (in the remainder of the text denoted by chemical repellents), the PHA/PHB neurons responded with short peaks in intracellular calcium.

### IFT components accumulate at the ciliary tip upon stimulation with a hyperosmotic solution

Next, we imaged the cilia, the organelles that protrude from the surface of the chemosensory neurons and require IFT^27^ (Figure 2A) for formation and maintenance. To visualize what happens in the cilia when the nematode tail is exposed to a chemosensory stimulus, we generated time-averaged image stacks and kymographs of fluorescently labeled IFT motors and particles in the phasmid cilia, before, during and after exposure to a hyperosmotic or chemical stimulus. For this purpose, we performed epi-fluorescence microscopy of live *C. elegans* strains that express fluorescently labeled kinesin-II (KAP-1::eGFP); OSM-3 (OSM-3::mCherry); IFT dynein (XBX-1::eGFP); IFT-particle subcomplex A (CHE-11::mCherry) or subcomplex B (OSM-6::eGFP), in the microfluidics chip.

**Figure 2:**
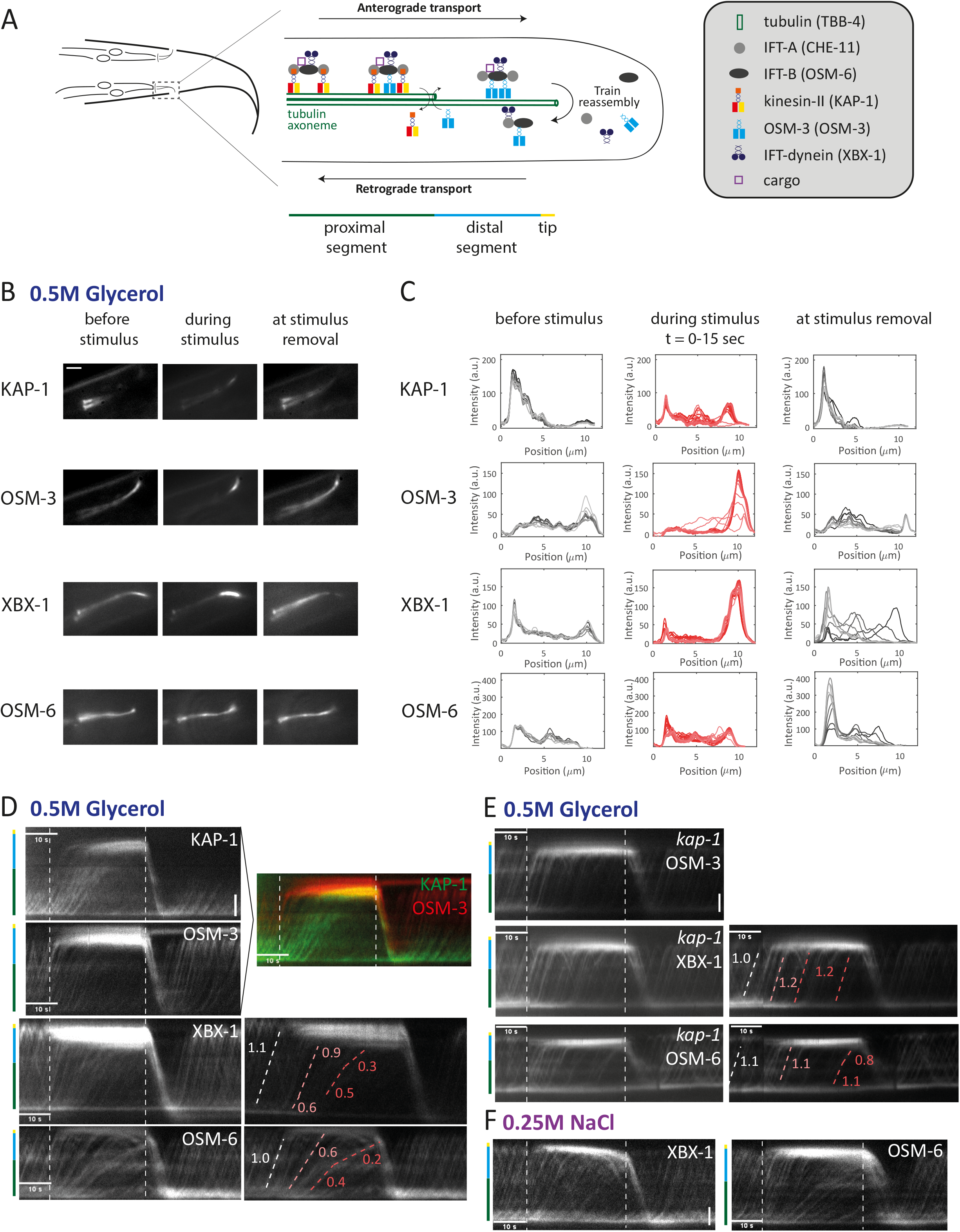
In response to hyperosmotic stimuli, IFT components accumulate at the ciliary tip due to the inhibition of retrograde transport. See also Movie S4, Figure S5, S6 and S7. A Schematic overview of the chemosensory cilia and intraflagellar transport (IFT), including key components. In the remainder of the figure, the proximal segment is shown in green, the distal segment in blue and the tip in yellow, as depicted. B Fluorescence intensity images time-averaged over 4.5 seconds of KAP-1::eGFP (kinesin-II), OSM-3::mCherry, XBX-1::eGFP (IFT dynein) and OSM-6::eGFP (IFT-B), before the start of the stimulus (left images), during glycerol exposure (middle images), after removal of the stimulus (right images). Scale bar: 2 µm. C Fluorescence intensity as a function of position, plotted at different times. Traces generated from the kymographs shown in 2D. Black curves correspond to intensities before glycerol exposure, red curves during exposure and grey curves after stimulus removal. See Figure S5A for 3D plot of this data set. D Kymographs (space-time plots) of KAP-1::eGFP (kinesin-II), n=7; OSM-3::mCherry, n=10; XBX-1::eGFP (IFT dynein), n=13 and OSM-6::eGFP (IFT-B), n=10; upon glycerol exposure. Vertical: position in cilium (color bar left indicates ciliary compartment), scale bar: 2 µm; horizontal: time. Dotted line left indicates the start of the stimulus, dotted line right stimulus removal. Representative kymographs of n=x animals. For XBX-1 and OSM-6 velocities (µm/s) were quantified, as shown on the right. E Kymographs of OSM-3::mCherry, n=11; XBX-1::eGFP (IFT dynein), n=11; and OSM-6::eGFP (IFT-B), n=8; in kap-1 mutant background upon glycerol exposure. For XBX-1 and OSM-6 velocities (µm/s) were quantified, as shown on the right. F Kymographs of XBX-1::eGFP, n=9; and OSM-6::eGFP, n=12; upon exposure to 0.25M NaCl.

Before stimulation with a chemosensory trigger, IFT dynein and IFT-B are distributed throughout the proximal and distal segments, with accumulations at the ciliary base (Figure 2B). Kinesin-II, the import motor that drives transport in the proximal segment of the cilium, also shows accumulation at the base and is mostly found at the proximal segment. OSM-3, the second anterograde motor that gradually takes over transport from kinesin-II in a region near the end of the proximal segment, is mostly found in the distal segment. These observations are in line with previous reports.^21,24,28^ During stimulation with glycerol, the movies (Movie S4) and time-averaged fluorescence images (Figure 2B) of all IFT motors and components reveal increased fluorescence intensities at the ciliary tip, indicating redistribution and accumulation of the IFT machinery at the tip.

We had a closer look at this redistribution by plotting the average fluorescence intensity along the cilium before, during and after glycerol stimulation for kinesin-II, OSM-3, IFT dynein and IFT-B (Figure 2C and Figure S5A). For OSM-3 and IFT dynein we observed increased fluorescence intensities at the ciliary tip. IFT-B also shows increased fluorescence intensity at the tip, although, to a lesser extent than IFT dynein and OSM-3. Very remarkably, kinesin-II, which is normally mostly present in the proximal segment, shows increased fluorescence intensity in the distal segment and tip, indicating unusual redistribution from the proximal towards the distal segment and tip. After stimulus removal, the fluorescence intensities at the tip decrease and those at the base increase, indicating a quick redistribution of the motors and components away from the ciliary tip.

To obtain further insight in the dynamics of the redistribution of the IFT machinery we used Kymograph Clear^29^ to generate kymographs of kinesin-II, OSM-3, IFT-dynein and IFT-B movement along the cilium before, during and after stimulation with glycerol (Figure 2D). In line with the time-averaged fluorescence images and intensity profiles, the kymographs show an increased fluorescence intensity at the ciliary tip for all IFT motors and components upon stimulation with glycerol, indicating accumulation at the tip. Based on these observations, we wondered which of the IFT particles accumulates at the tip first. Because we used a worm strain expressing both fluorescently labeled kinesin-II and OSM-3, we could visualize the behavior of both anterograde motors in the same animal. The kymograph reveals that OSM-3, the anterograde motor that mostly drives transport in the distal segment of the cilium, starts accumulating at the tip first. Indeed, to our big surprise, as we already observed in the time-averaged fluorescence images and intensity profiles, kinesin-II subsequently moves all the way into the distal segment. It continues transporting anterograde trains for much longer during the stimulus than OSM-3 and at a certain moment it even becomes the sole functional anterograde motor driving transport along the entire axoneme. Eventually, also kinesin-II is completely accumulated at the tip and depleted from the rest of the cilium. The kymograph overlay of kinesin-II and OSM-3 clearly shows the order of events, with OSM-3 accumulating at the tip first, followed by kinesin-II moving into the distal segment, and finally kinesin-II accumulating at the tip. The retrograde Fourier-filtered kymograph of OSM-3 shows that retrograde transport of OSM-3 stops immediately at the start of the stimulus, which might explain the accumulation at the tip (Figure S5B).

Next, we had a closer look at IFT dynein, the retrograde motor. IFT dynein showed an increase in the fluorescence intensity at the tip and a decrease at the base (Figure 2D), which indicates that IFT dynein redistributes from base to tip, immediately after stimulus delivery. The anterograde and retrograde Fourier-filtered kymographs of IFT dynein indicate that, immediately after the start of the stimulus, retrograde transport stops. Since IFT dynein drives all retrograde IFT, this must be the cause of the cease of retrograde OSM-3 transport indicated above. On the other hand, anterograde transport of IFT dynein continues for a while after stimulus delivery, seemingly as long as the “reservoir” of IFT dynein at the base is not depleted (Figure S5B). During the stimulus exposure, the slopes of the kymograph lines become less steep, indicating that anterograde transport slows down. We observed a slow down in anterograde transport for all animals measured. We determined the IFT-dynein velocities by calculating the slope of the lines in the kymographs in Figure 2D. Before stimulus exposure we measured a velocity of 1.1 µm/s; during the glycerol exposure the velocity initially decreased to 0.6 µm/s in the proximal and 0.9 µm/s in the distal segment, and eventually decreased to 0.5 µm/s and 0.3 µm/s, respectively (Figure 2D). Also, IFT subcomplex B accumulates at the tip, while its “reservoir” at the base decreases during the stimulation (Figure 2D). Transport continues during the stimulus, and as for IFT dynein, the velocity decreases, from 1.0 µm/s before stimulus exposure to 0.6 µm/s during glycerol exposure to eventually a velocity of 0.4 µm/s in the proximal and 0.2 µm/s in the distal segment (Figure 2D). The initial slowdown can be explained by (slower) kinesin-II taking over the role of (faster) OSM-3 in most of the cilium, including the distal segment. In contrast to IFT dynein however, IFT-B appears to be transported from base to tip for the entire duration of the stimulus. The kymograph lines become increasingly less steep, indicating that IFT-B slows down even further than can be expected on basis of kinesin-II^30^ taking over from OSM-3. The further slowdown might be due to only a very small number of kinesin-II motors being attached to each train, too few to be discernable in the kinesin-II kymographs, that only intermittently drive transport of the trains. IFT subcomplex A behaves in a similar way as IFT-B (Figure S6).

We noticed that anterograde transport of OSM-3 appears to stop earlier than that of IFT dynein and IFT-B, demonstrating that OSM-3 reaches a new equilibrium during the glycerol stimulus sooner. An explanation for this could be that, at the base, excess IFT dynein and IFT-B appear to be present in a “reservoir”. As long as the reservoir has not been depleted, anterograde transport can continue. No such reservoir of OSM-3 seems to exist, which can explain why the anterograde transport of OSM-3 appears to stop earlier.

We hypothesized that the decreased velocity of anterograde trains was caused by kinesin-II, which has a ∼2.5x lower velocity than OSM-3,^30^ acting as sole anterograde motor. To test our hypothesis, we performed a glycerol stimulus to worms lacking kinesin-II function (*kap-1*), in which OSM-3 drives anterograde transport over the entire length of the cilium.^30^ In response to glycerol, we observed a similar response in *kap-1* worms as in wildtype worms, with OSM-3, IFT dynein and IFT-B accumulating at the tip (Figure 2E). In contrast to wildtype worms, however, the trains did not slow down. For XBX-1 we estimated the velocity to be 1.0 µm/s before and 1.2 µm/s during glycerol exposure; for OSM-6 we estimate a velocity of 1.1 µm/s before stimulus exposure and 0.8-1.1 µm/s during glycerol exposure (Figure 2E). This confirms that anterograde IFT slow down in wild-type worms is caused by kinesin-II acting as the sole anterograde motor.

Upon removal of the hyperosmotic stimulus, densely packed retrograde trains appear in the kymographs and the fluorescence intensity at the base increases (Figure 2D), caused by a sudden flux of all IFT motors and particles back to the base. Subsequently, within about 10 seconds, IFT dynamics reestablishes similar to the situation before the stimulus. Remarkably, this recovery of pre-stimulus IFT occurs more quickly than tip accumulation following stimulus exposure. An explanation for this could be that IFT-dynein-driven retrograde transport (causing recovery) is ∼2.5 times faster than kinesin-II-driven anterograde transport (causing tip accumulation).

In response to high concentrations of NaCl we observed a similar reaction of the IFT machinery as to glycerol (Figure 2F and Figure S7).

Taken together, our results show that exposure to hyperosmotic stimuli leads to accumulation of the IFT machinery at the ciliary tip, likely due to inhibition of retrograde trains departing from the tip. Upon stimulus removal, tip-accumulated IFT components return back to the base in a wave, after which IFT quickly restores to the situation prior to stimulus exposure.

### IFT components redistribute towards the ciliary base upon stimulation with chemical repellents

When we exposed the worm tail to SDS, we observed a different response of the ciliary IFT machinery. The time-averaged images after 75 seconds SDS exposure (Figure 3A) show that the intensities of the IFT motors and particles at the ciliary tip have decreased. For OSM-3, IFT dynein and IFT-B some fluorescence can still be detected in the proximal segment, while kinesin-II is only detectable at the ciliary base. In contrast to exposure to a hyperosmotic stimulus, which leads to accumulation of the IFT machinery at the tip, these observations indicate that exposure to SDS leads to redistribution of the IFT machinery away from the tip and distal segment towards the base.

**Figure 3:**
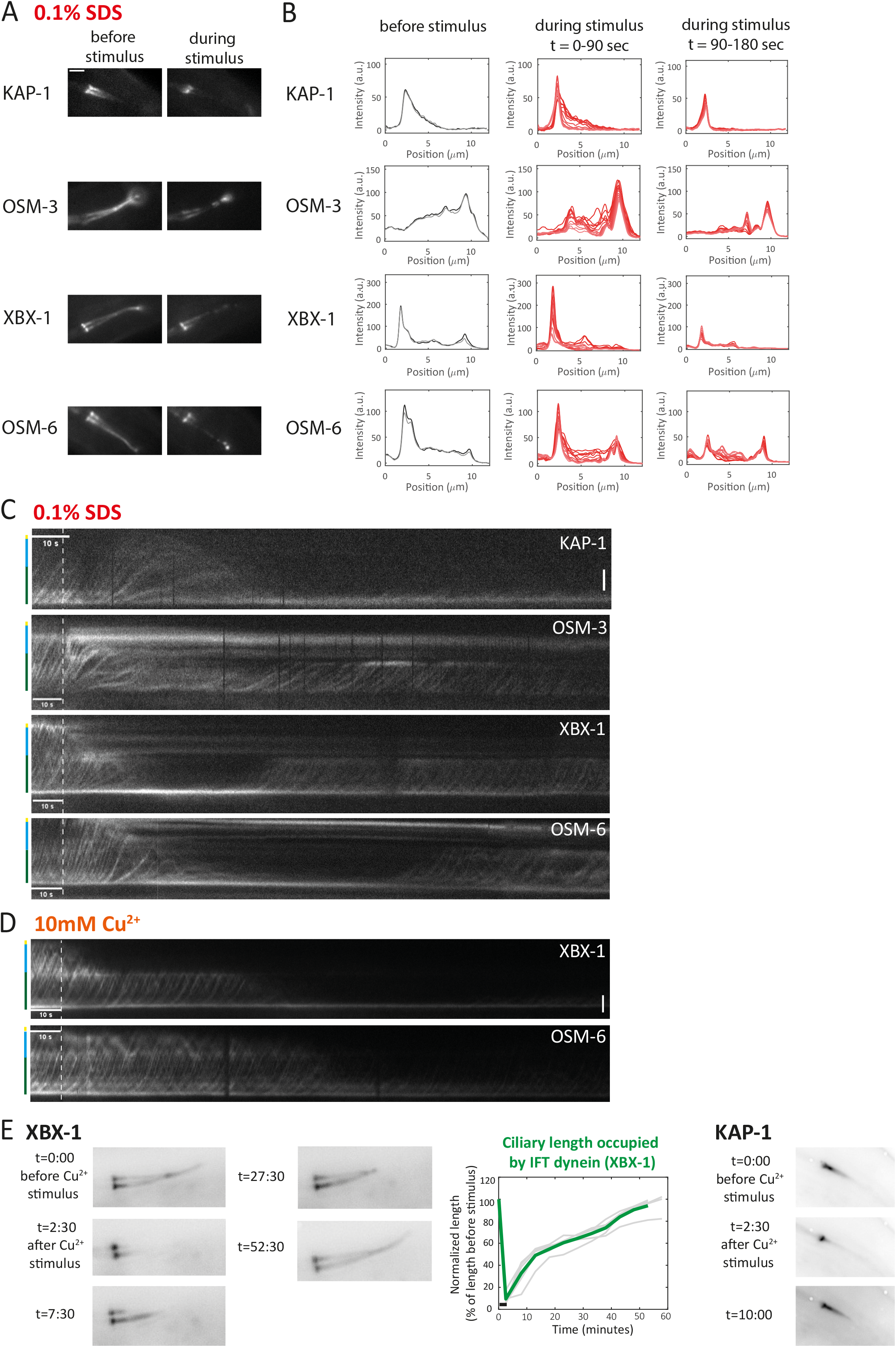
In response to chemical repellents SDS or Cu^2+^, the ciliary axoneme shrinks and IFT components redistribute towards the ciliary base. See also Figure S6 and S8. A Fluorescence intensity images time-averaged over 4.5 seconds of KAP-1::eGFP (kinesin-II), OSM-3::mCherry, XBX-1::eGFP (IFT dynein) and OSM-6::eGFP (IFT-B), before the start of the stimulus (left images) and 75 seconds after the start of the SDS exposure (right images). Scale bar: 2 µm. B Fluorescence intensity as a function of position in the cilium plotted for different time poins. Traces generated from the kymographs shown in 3C. Black curves correspond to intensities before, red curves during SDS exposure. See Figure S8A for 3D plot of this data set. C Kymographs of KAP-1::eGFP (kinesin-II), n=9; OSM-3::mCherry, n=11; XBX-1::eGFP (IFT dynein), n=16 and OSM-6::eGFP (IFT-B), n=8; before and during SDS exposure. Vertical: position in cilium (color bar indicates ciliary component), scale bar: 2 µm; horizontal: time. The dotted vertical line indicates the start of the stimulus. The stimilus lasts for the entire remainder of the kymograph. Representative kymographs of n=x animals. D Kymographs of XBX-1::eGFP, n=11; and OSM-6::eGFP, n=8; before and during exposure to Cu^2+^. E Left: inverted fluorescence images of IFT dynein (XBX-1::eGFP) before and during a 2.5 minutes Cu^2+^ stimulus followed by exposure to M13 buffer. t = time in minutes:seconds. Graph represents the ciliary length occupied by IFT dynein before, during and after 2.5 minutes exposure to Cu^2+^ followed by exposure to M13 buffer. Light grey lines represent individual measurements on one cilium, thick green line the average. Black horizontal bar indicates the onset and duration of the stimulus. n=5 cilia of n=3 worms. Right: inverted fluorescence images of kinesin-II (KAP-1::eGFP) before, during and after 2.5 minutes Cu^2+^ stimulus followed by exposure to M13 buffer. t = time in minutes:seconds.

To obtain further insight in this redistribution, we generated plots of the average fluorescence intensity along the cilium before and during SDS stimulation for kinesin-II, OSM-3, IFT dynein and IFT-B (Figure 3B and Figure S8A). The plots show that a few seconds after the start of the SDS stimulus, the fluorescence intensities decrease at the tip and the distal segment and increase at the proximal segment and base. The decrease of fluorescence intensity at the tip is more pronounced for XBX-1 (for which fluorescence at the tip disappears almost completely), than for OSM-3 and IFT-B (for which some fluorescence remains at the tip).

To study the dynamics of the redistribution of the IFT machinery, we generated kymographs of kinesin-II, OSM-3, IFT dynein and IFT-B movement along the cilium before and during stimulation with SDS (Figure 3C). The kymographs indicate that the redistribution of the IFT components happens in two steps. First, the motors and particles retract from the distal segment and transport is only observed in the proximal segment. Second, also transport in the proximal segment stops and all components have redistributed to the base. The velocity of motors and particles starts to decrease about 10-20 seconds after the start of the stimulus until transport comes to a complete halt about 20-30 seconds later. For kinesin-II, IFT dynein and IFT-B, a clear increase in fluorescence intensity can be observed at the base. Remarkably, OSM-3, IFT dynein and IFT-B redistribute towards the base, but a substantial fraction of these proteins remains at the tip. After approximately 2 minutes stimulus exposure, IFT starts to recover to normal velocities, but only in the proximal segment. IFT-A behaved similarly as IFT-B (Figure S6).

Upon stimulation with Cu^2+^ a similar reaction of the IFT machinery is observed (Figure 3D and Figure S8B). The response to Cu^2+^ was, however, somewhat subtler than to SDS, which makes the two-step redistribution process (first from distal to proximal segment, followed by a complete redistribution towards the base) more clearly visible in the kymographs.

To test whether IFT-component distribution recovers and how long it takes, after switching from Cu^2+^-stimulus back to buffer, long-time imaging over up to an hour was required. Continuous imaging proved to be difficult because of photobleaching and movement of the (sedated) animal. To overcome these limitations we imaged the distribution of the IFT machinery with time intervals of 5 minutes, using IFT dynein as a marker. We observed in these images (Figure 3E) that recovery is a gradual process on the tens of minutes timescale and appears to be complete after about 55 minutes. Kinesin-II, which only spans the proximal segment of the cilium, appears to be recovered to its pre-stimulus distribution after approximately 10 minutes (Figure 3E).

In short, exposure to a chemical repellent, such as a solution of SDS or Cu^2+^, leads to redistribution of IFT motors and particles from the distal to the proximal segment and eventually all the way to the ciliary base. This process is reversible. After removal of the stimulus, IFT recovers on a timescale of about an hour. The response to a chemical repellent is in sharp contrast to the response to a hyperosmotic stimulus, which causes strong accumulation of IFT components at the ciliary tip.

### The ciliary axoneme shrinks in response to chemical stimuli, but the ciliary membrane stays intact

Because IFT is required for the maintenance and functioning of cilia, we wondered what the effects of exposure to hyperosmotic stimuli and chemical repellents would be on ciliary structure. We therefore performed fluorescence microscopy of *C. elegans* strains expressing fluorescently labeled tubulin (TBB-4::eGFP). Because tubulin is the subunit of the microtubule-based axoneme, this worm strain provides insight in the structure of the axoneme during stimulus exposure. In addition to tubulin, we imaged the transmembrane calcium channel OCR-2 (OCR-2::eGFP). Because OCR-2 is a transmembrane protein abundantly expressed in cilia (with a substantial accumulation at the tip),^31,32^ this provides insight in the shape of the ciliary membrane during stimulus exposure. Moreover, OCR-2 is known to play a crucial role in the downstream signaling of G protein coupled receptor (GPCR) activation and hyperosmotic stimuli are known to directly activate OCR-2 channel opening.^3^ We have previously shown that OCR-2 is partially transported via IFT and partially undergoes diffusion.^32^

In response to hyperosmotic triggers, IFT motors and particles accumulate at the tip of the cilium. We therefore expected the microtubule-based axoneme to stay intact and indeed, the time-averaged fluorescence images and kymographs of tubulin do not show any changes upon exposure to glycerol (Figure 4A,B). The time-averaged fluorescence images and kymograph of OCR-2 indicate that the concentration of OCR-2 increases at the tip upon stimulation with glycerol (Figure 4A,B). By quantifying the fluorescence intensity of OCR-2::eGFP specifically at the ciliary tip, we indeed observed a substantial increase of OCR-2 upon stimulation with glycerol (Figure 4C). Upon removal of the stimulus, a wave of OCR-2 molecules away from the tip, back towards the distal and proximal segment can be observed (Figure 4A-C). Most likely, these molecules are connected to IFT trains at the moment of stimulus removal and are transported back, driven by IFT dynein. A fraction of OCR-2 remains at the tip, suggesting that these molecules were not connected to IFT trains at the moment of stimulus removal. This result shows that the ciliary membrane retained a similar shape in response to exposure to a hyperosmotic stimulus and that the calcium channel OCR-2 is further enriched at the tip.

**Figure 4:**
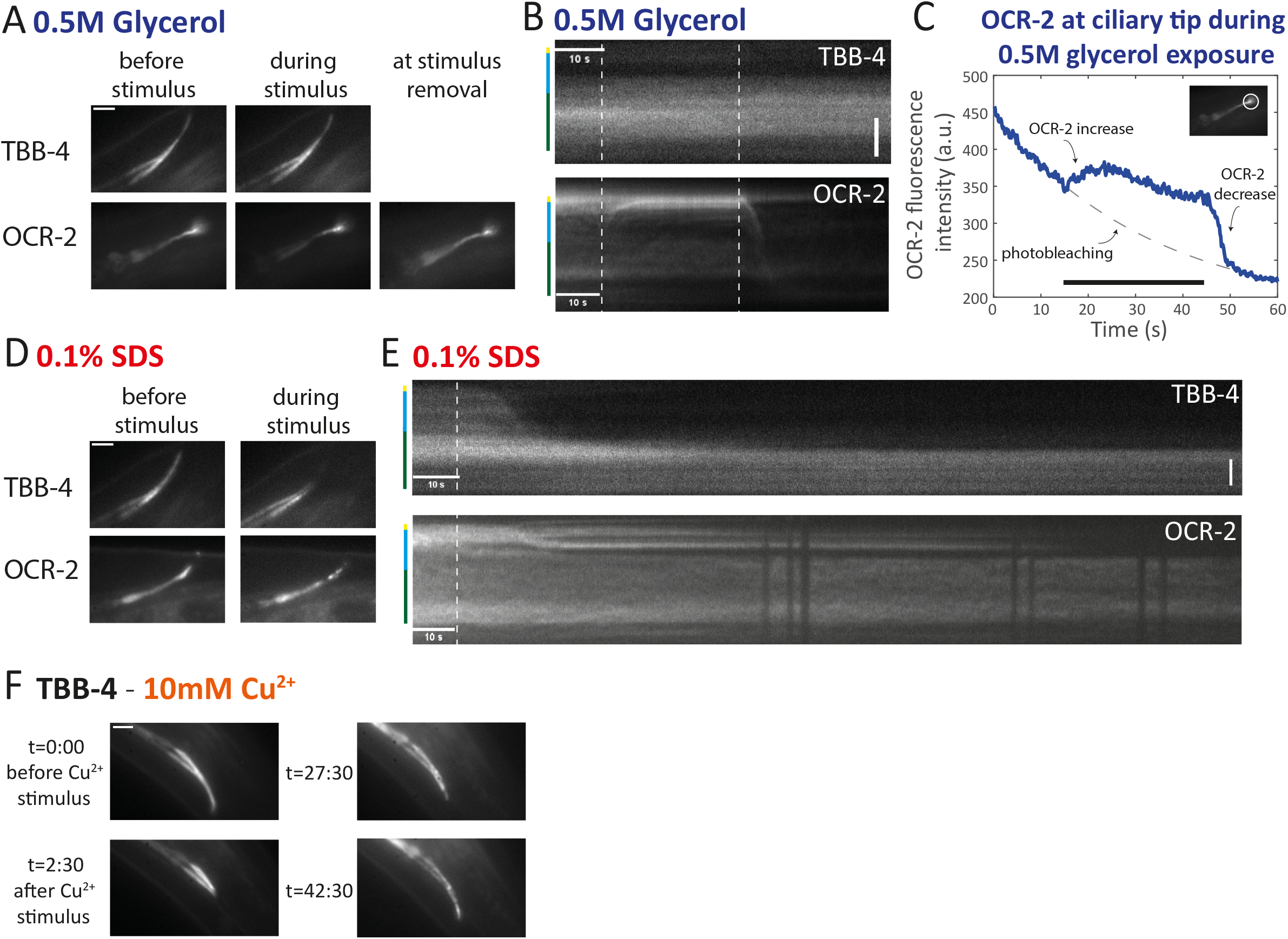
Structure and function of the cilium in response to hyperosmotic and chemical stimuli. A Fluorescence intensity images time-averaged over 4.5 seconds of TBB-4::eGFP and OCR-2::eGFP before (left images) and during glycerol exposure (middle images) and after stimulus removal (right image). Scale bar: 2 µm. B Kymographs of TBB-4::eGFP and OCR-2::eGFP during and after glycerol exposure. Vertical: position in cilium (color bar indicates ciliary component), scale bar: 2 µm; horizontal: time. First dotted line indicates the start of the stimulus, second dotted line end of stimulus. TBB-4, n=4; OCR-2, n=12. Representative kymographs of n=x animals. C Time trace of the fluorescence intensity of OCR-2::eGFP (y-axis) at the ciliary tip (indicated by the white circle in the insert image) plotted over time. Black horizontal bar indicates the onset and duration of 0.5M glycerol stimulus. Grey dotted line is an exponential fit representing the decay of the fluorescence intensity due to photobleaching. D Fluorescence intensity images time-averaged over 4.5 seconds, of TBB-4::eGFP and OCR-2::eGFP before the stimulus (left images) and 75 seconds after the start of the SDS exposure (right images). Scale bar: 2 µm. E Kymographs of TBB-4::eGFP and OCR-2::eGFP during and after SDS exposure. Vertical: position in cilium (color bar indicates ciliary component); horizontal: time. Dotted line indicates the start of the stimulus. TBB-4, n=9; OCR-2, n=12. Representative kymographs of n=x animals. F Inverted fluorescence images of tubulin (TBB-4::eGFP) before and after 2.5 minutes Cu^2+^ stimulus (exposure to M13 buffer after the stimulus). t = time indicated in minutes:seconds. Scale bar: 2 µm.

In response to the chemical repellent SDS, the cilium reacts very differently. Roughly 10 seconds after the start of SDS exposure, tubulin redistributes from the distal segment to the proximal segment (Figure 4D,E), indicating axonemal shortening of the singlet microtubules. We hypothesize that this is a direct consequence of the IFT machinery redistributing out of the distal segment immediately following SDS exposure (Figure 3A). This, in turn, might prevent new tubulin subunits from being incorporated at the microtubule plus ends, resulting in catastrophes of the singlet microtubules. The kymograph indicates that the proximal segments remain intact and no further shortening occurs, presumably because the doublet microtubules in the proximal segment are more stable than the singlet microtubules in the distal segment.^33^ The transmembrane protein OCR-2 partially redistributes from the tip into the distal segment (Figure 4D,E). A fraction of OCR-2 remains at the tip, suggesting that the ciliary membrane is not remodeled, e.g. by “following” the shortening of the axoneme. Eventually also these molecules disappear from the tip, but not to the same extent as tubulin, indicating that even though the axoneme shortens, the ciliary membrane remains.

To test whether the shortening of the axoneme in response to SDS (Figure 4D,E) and Cu^2+^ (Figure S8B) is reversible, we imaged the TBB-4 distribution with time lags. As already shown in the kymographs, the distal segments disappeared after 2.5 minutes of Cu^2+^ exposure, while the proximal segments stayed intact. Imaging at different time points after the removal of the stimulus revealed that the distal segments recovered (Figure 4F) at an approximately similar time scale as the recovery of IFT (Figure 3E).

In summary, in response to a hyperosmotic solution the ciliary membrane stays at its place and there are no overall changes in the structure of the axoneme. Hyperosmotic exposure does lead to redistribution of the calcium channel OCR-2 towards the tip. Exposure to the chemical repellent SDS, reveals shortening of the axoneme until only the proximal segment remains. The transmembrane protein OCR-2 only partially redistributes away from the tip, indicating that the ciliary membrane does not retract.

### Intact axoneme and functioning IFT are required for calcium spiking

We wondered whether the nematode would still be able to sense a chemical trigger leading to neuronal activation, during the redistribution of all IFT motors and particles to the base and shortening of the ciliary axoneme in response to SDS and Cu^2+^ (Figure 3). We therefore stimulated GCaMP6s-expressing worms for 2.5 minutes with a chemical repellent, SDS or Cu^2+^, to cause shortening of the axoneme and redistribution of IFT components to the base. Thereafter, we briefly switched to buffer and after about 30 seconds gave a second stimulus. At this point, no calcium spiking was observed (Figure 5A). We did not observe a GCaMP6s signal in the cilium and dendrite nor an increased GCaMP6s signal in the soma, highlighting the crucial role of the cilium in sensing the chemical environment and evoking calcium spiking. In contrast, using glycerol as a stimulus (which we have shown to result in accumulation of the IFT machinery at the tip (Figure 2) and no axonemal shrinking (Figure 4)), we did observe a calcium spike in the somas of the chemosensory neurons upon a second stimulus (Figure 5B). We next tested whether the neuron would regain the ability to generate calcium spikes after 55 minutes of exposure to buffer following a first stimulus with Cu^2+^. This should be enough time for the IFT-component distribution to recover to the pre-stimulus situation. Indeed, a clear GCaMP6s signal could be observed in the dendrite and soma, qualitatively similar to the reaction to the first stimulus (Figure 5C), which confirms that a recovered cilium is able to sense the chemical environment resulting in calcium spikes.

**Figure 5:**
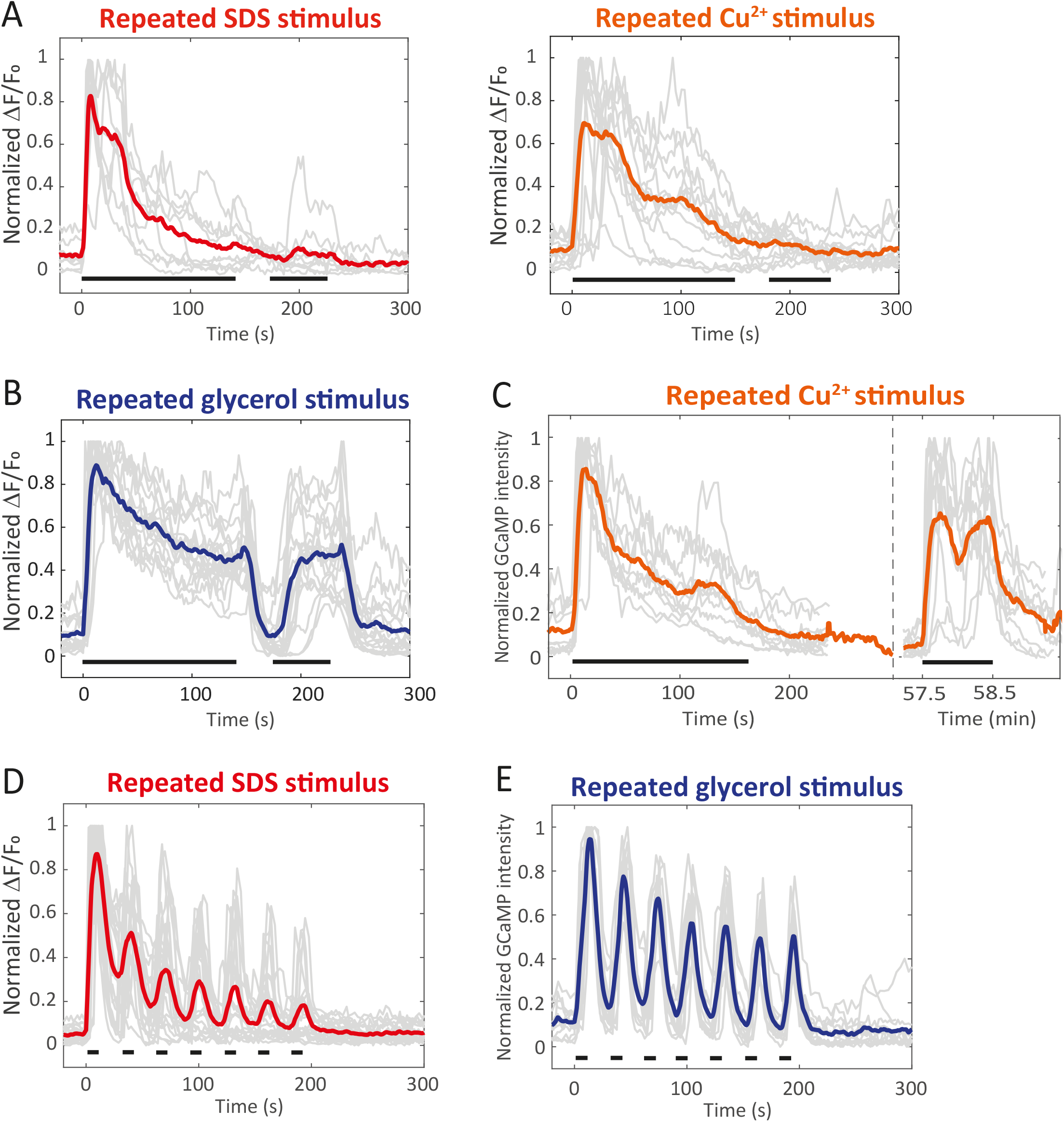
*C. elegans* habituates to a chemical repellent, but not to a hyperosmotic solution. A-E Graphs represent normalized GCaMP6s fluorescence intensity time trace in the PHA/PHB neurons in response to the stimuli as indicated below. Light grey lines are individual measurements, thick colored lines averages. Black horizontal bars indicate onset and duration of the stimulus. n=x-y, in which x indicates the number of somas and y the number of animals (see Methods). A 2.5 minutes exposure to 0.1% SDS, n=9-8 (left) and 10mM Cu^2+^, n=14-8 (right), followed by 0.5 minutes M13 buffer and a subsequent additional 1 minute exposure to the earlier stimulus. B 2.5 minutes exposure to 1M glycerol, followed by 0.5 minutes M13 buffer and a subsequent additional 1 minute exposure to glycerol. n=19-8 C 2.5 minutes exposure to 10mM Cu^2+^, followed by 55 minutes M13 buffer (indicated by dashed line) to allow full recovery of axoneme and IFT and subsequent 1 minute exposure to 10mM Cu^2+^. n=10-6 D-E Repeated stimuli of 10 seconds of 0.1% SDS, n=22-12 (D) and 1M glycerol, n=13-10 (E) alternated with 20 seconds M13 buffer.

Taken together, these results indicate that after prolonged chemical stimulation with SDS and Cu^2+^ the neuron shows habituation. Habituation is the adaptation of the sensory system to an external signal by evoking lower neuronal activity upon subsequent exposures. Remarkably, hyperosmotic stimuli do not appear to provoke habituation. To further confirm this, we tested repeated short exposures to SDS alternated with buffer. The GCaMP6s fluorescence response became lower upon every new exposure, till it was almost negligible the seventh time (Figure 5D). When we performed repeated stimuli with glycerol, only a slight decay in GCaMP6s fluorescence intensity was observed, similar to long, continuous glycerol exposure (Figure S3B). The neurons clearly kept responding to glycerol (Figure 5E), indicating that the habituation that we observed for SDS, did not happen for a hyperosmotic stimulus.

## Discussion

*C. elegans* is well-known to perform avoidance behavior when exposed to repellent chemicals.^3^ In the current study we have made use of a microfluidics chip to apply stimuli with a high degree of temporal control. This allowed studying the multiscale process of chemosensation: neuronal activity, which eventually leads to behavioral changes, as well as the response of the ciliary IFT machinery. We found that hyperosmotic solutions and chemical repellents resulted in distinct activities of the phasmid chemosensory neurons. In response to hyperosmotic solutions, the PHA/PHB neurons reacted with increased intracellular calcium concentrations for the entire duration of the stimulus, whereas chemical repellents caused short peaks in intracellular calcium concentration.

In a previous study^34^ it was shown that the ASH neuron, located in the head of the worm, shows similar calcium traces in response to solutions of Cu^2+^ and glycerol as the PHA/PHB neurons that we describe in the current study. Exposing a worm to 2 and 10 mM of Cu^2+^ led to sharp peaks in intracellular calcium in the ASH neuron,^34^ while stimulation with 0.5M glycerol caused a prolonged increase in intracellular calcium, for as long as the stimulus lasted.^17,34^ Taking together our data and these previous observations, the distinct calcium traces in response to hyperosmotic and chemical stimuli appear not to be limited to the phasmid neurons only.

It should be kept in mind that the worms were anesthesized using levamisole for 10 minutes prior to exposing them to a chemosensory stimulus. Although the phasmid neurons are not cholinergic, they innervate cholinergic neurons.^35^ We can therefore not completely exclude that levamisole has an indirect effect on the activity of the phasmid neurons.

Eukaryotic cells have been shown to regulate intracellular calcium levels in response to changes in osmotic pressure. When exposed to hyperosmotic stress, the intracellular calcium concentration increases in order to equalize the osmotic pressure on both sides of the membrane.^36^ It might be that the calcium spike that we observed in response to glycerol and high salt is not primarily a neuronal spike, but a mechanism to cope with hyperosmotic stress. This could explain why GCaMP6s fluorescence intensity stayed high for the entire duration of the stimulus.

In a recent study, the avoidance behavior of *C. elegans* was investigated in response to an upshift of osmolarity.^37^ It was reported that *C. elegans* shows gradually increasing reorienting maneuvers in response to increasing osmolarity, different from acute escape-like avoidance behavior. The slow aversion to osmotic upshift was shown to require the cGMP-gated channel TAX-2. Taken together with our results, this suggests that both calcium spiking as well as avoidance behavior is different in response to hyperosmotic solutions or chemical repellents. It remains to be investigated if the typical avoidance behavior that is caused by SDS or Cu^2+^ is regulated via a different signaling pathway than the aversive behavior towards hyperosmotic solutions.

Not only the neuronal responses to hyperosmotic solutions were different from chemical repellents, also the response of the IFT machinery was completely different. While chemical stimuli caused the IFT machinery to redistribute towards the ciliary base, hyperosmotic stimuli resulted in a redistribution towards the tip. This tip accumulation of IFT components upon exposure to a hyperosmotic solution appears to be caused by an inhibition of retrograde transport. The inhibition specifically takes place at the ciliary tip, since some retrograde trains could still be observed in the proximal segment at the start of the stimulus (Figure S5B). Under normal conditions, anterograde trains arrive at the ciliary tip, disassemble and reassemble before taking off as IFT-dynein driven retrograde trains, back to the base.^38^ Hyperosmotic stimuli appear to specifically prevent this process, by either disrupting the train disassembly / reassembly process, or by preventing activation of IFT dynein, from an inactive cargo on anterograde trains to the active motor on retrograde trains. Further experiments will be required to shed light on this, for example by tracking the behavior of individual IFT components around the tip.^38^

Next to the inhibition of retrograde transport, a striking observation following hyperosmotic stimulation was that kinesin-II moves all the way into the distal segment and accumulates at the tip. In the normal situation, without stimulus, kinesin-II cycles between ciliary base and the proximal segment, acting as anterograde motor for IFT trains entering the cilium. In the proximal segment, kinesin-II stochastically detaches from anterograde trains, reattaching to retrograde trains in order to be transported back to the base.^22,24^ It has been observed before that in worms deficient of MAP kinase DYF-5, kinesin-II enters the distal segment of the cilium.^39^ From this data it was inferred that DYF-5 could play a role in the undocking of kinesin-II from IFT particles and the docking of OSM-3 onto IFT particles.^39^ It could be that, in our experiments, hyperosmotic stress has an effect, direct or indirect, on the function of DYF-5, causing changes in motor regulation. If DYF-5 activity were required for the undocking of kinesin-II from IFT particles, inhibition of DYF-5 could explain movement of kinesin-II into the distal segment. Another explanation of the effect of hyperosmotic stimuli on kinesin-II distribution could be that this is a direct consequence of the termination of retrograde trains leaving from the ciliary tip. At some point after the stimulus, kinesin-II motors detached from anterograde trains will not encounter retrograde trains anymore to transport them back to the base. Instead, they will only encounter other anterograde trains, to which they might dock and be transported further along the cilium towards the tip. Such undocking from and docking to different anterograde trains has recently been observed in single-molecule experiments.

Exposure to the chemical stimuli SDS and Cu^2+^ led to redistribution of IFT components towards the ciliary base and disappearance of the distal part of the axoneme. This shortening of the axoneme appeared to be slower than the redistribution of the IFT machinery towards the base of the cilium, indicating that the collapse of the axoneme is a consequence of the absence of transport to the tip. Before, we have performed dendritic laser ablation to stop the entry of new material to the cilium.^33^ In those experiments we also observed redistribution of IFT motors and particles towards the base, which subsequently caused shortening of the axoneme. We furthermore showed that, in response to a chemical stimulus, both axonemal shrinkage and redistribution of the IFT machinery are reversible (Figure 3E and 4E). After removal of the stimulus, kinesin-II returned to the pre-stimulus situation in only about 10 minutes. This relatively short recovery is most likely caused by kinesin-II only being present in the proximal segment of the cilium, which stayed intact during stimulus exposure. In contrast, IFT recovered over the entire length of the cilium in about 55 minutes. These data show that it takes time, roughly one hour, to completely rebuild the distal segment after axonemal shrinkage caused by exposure to a chemical repellent.

We also looked at the response of transmembrane proteins involved in signaling following stimulation. Upon exposure to a hyperosmotic stimulus, we observed a redistribution of the calcium channel OCR-2 towards the ciliary tip (Figure 4A,B). We have shown before that OCR-2 is distributed over the cilium by a location-dependent interplay between IFT and diffusion within the membrane.^32^ The extra accumulation of OCR-2 upon hyperosmotic stimulation is thus most likely a direct consequence of the accumulation of the IFT machinery at the ciliary tip. It could be that an increase of OCR-2 at the ciliary tip plays a role in regulating the calcium response to hyperosmotic stress. It has been shown before that the calcium channel OCR-2 can be directly opened by hyperosmotic stimuli.^3^ It remains to be investigated whether receptors and other ion channels also accumulate at the ciliary tip. It would be interesting to find out whether transport of these proteins is actively regulated in response to stimuli, e.g. by affecting binding to the IFT machinery.

In response to a chemical stimulus, we observed partial redistribution of OCR-2 away from the ciliary tip (Figure 4C,D). Since OCR-2 is involved in the downstream signaling of GPCRs,^3^ fewer OCR-2 molecules at the tip could result in desensitization, i.e. habituation. Indeed, repeated exposure to a chemical repellent rendered the cilium incapable of sensing the stimulus when the axoneme was at its shortest and all IFT components were redistributed towards the base. When exposing the worm to a chemical stimulus repeatedly, with only short time between the stimuli, the calcium spikes decreased, pointing towards a mechanism of habituation. Previously, a similar adaptation to soluble repellents has been reported for the amphid chemosensory neuron ASH in response to Cu^2+^.^8^ That study showed that the calcium response in the ASH soma decreased on a time scale of tens of seconds during successive short exposures, causing behavioral adaptation.

Taken together, our study highlights the flexibility of ciliary structure and dynamics. The distinct responses to hyperosmotic solutions on the one hand and chemical repellents on the other reveal that strikingly different reactions of the IFT machinery eventually lead to a similar behavioral response, namely avoidance. How the changes in IFT affect calcium spiking or vice versa remains to be investigated, for example using indicators for other secondary messengers, such as cGMP,^40^ and worm strains in which key players of the signaling cascade have been knocked out. In short, our findings highlight the intricacy of the cilium as sensing hub: the cilium is able to sense and respond to different external cues in distinct ways resulting in coherent behavioral output.

## Acknowledgements

We thank Aniruddha Mitra for helpful discussions and support. We acknowledge financial support from the European Research Council under the European Union’s Horizon 2020 research and innovation programme (Grant agreement no. 788363; “HITSCIL”).

## Author contributions

Conceptualization and Methodology, C.W.B. and E.J.G.P.; Investigation, C.W.B. and G.H., Formal analysis, C.W.B., G.H. and N.D.; Writing – Original Draft, C.W.B. and E.J.G.P.; Writing – Review & Editing, C.W.B., G.H., N.D., J.v.K. and E.J.G.P.; Funding acquisition and supervision, E.J.G.P.

## STAR Methods

### Maintenance of worm strains

*C. elegans* strains were maintained according to standard procedure, on NGM plates, seeded with HB101 *E. coli*, at 20°C. In all experiments, young adult worms were used. All IFT reporter strains used in this study carry integrated single-copy transgenes, previously generated in our lab via MosSCI.^41^

### Worm strains

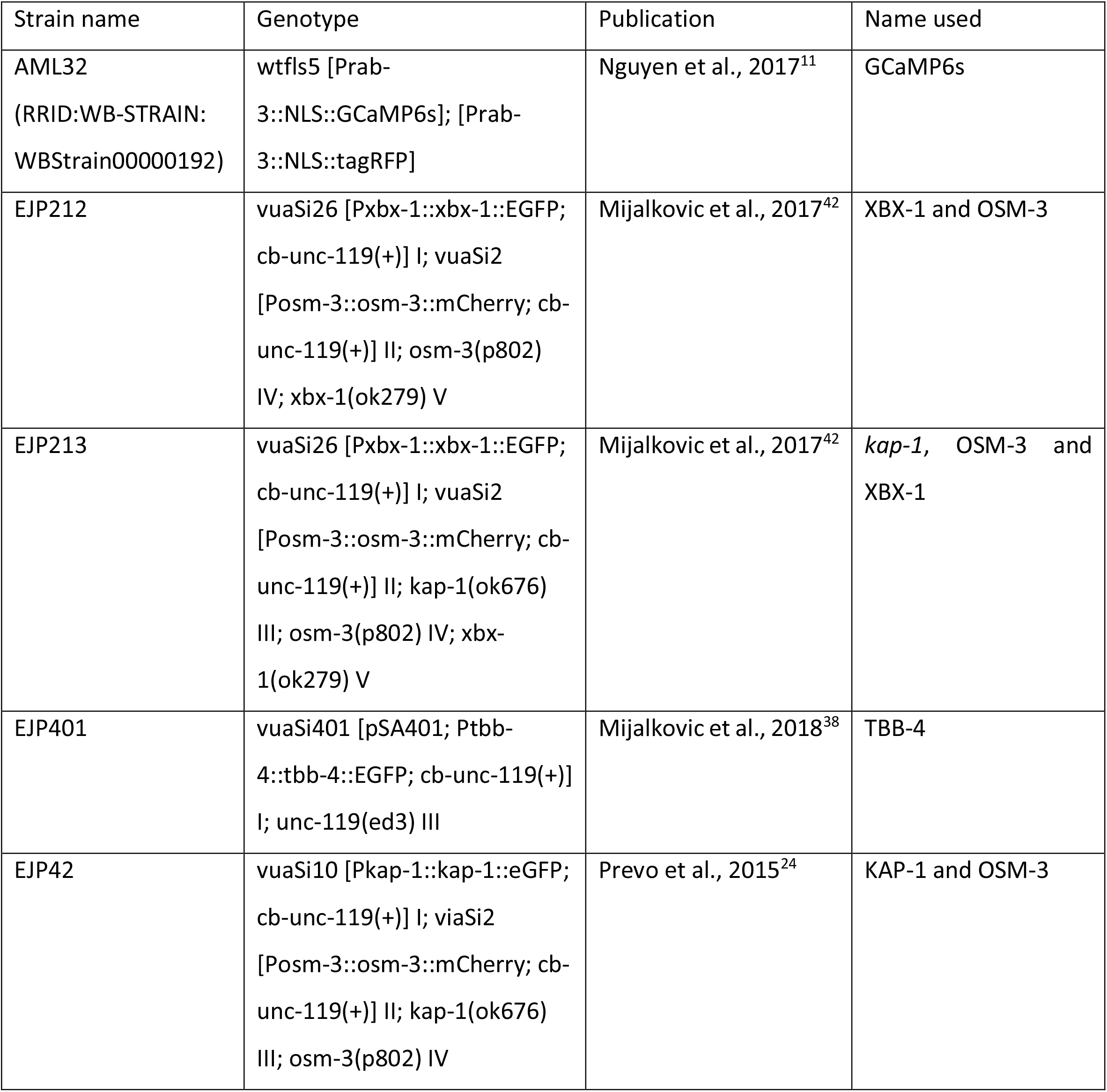
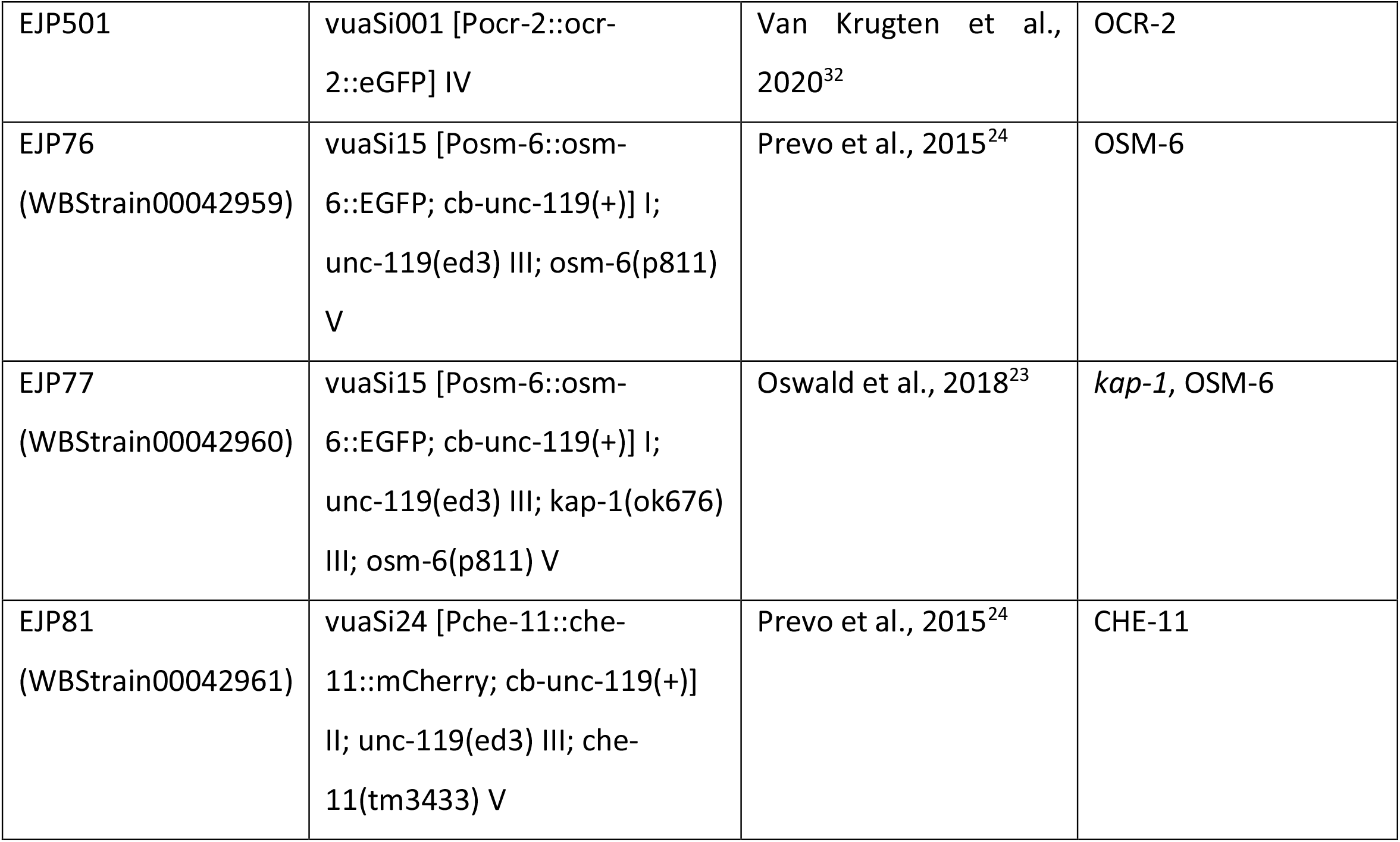

### Microfluidics chip design

The design of the microfluidics chip used in this study, was first presented by Chronis et al. (2007).^17^ A master with the desired pattern, consisting of a 26-32 μm layer SU-8-3025 photoresist on a silicon wafer, was made by AMOLF NanoLab Amsterdam. This master was repeatedly used to pour polydimethylsiloxane (PDMS) chips.

### Pouring PDMS chips

The two components of PDMS (Sylgard 184, Dow Corning) were mixed in a ratio of 1:10. After thoroughly mixing for at least five minutes, air bubbles were removed from the mixture by evacuating it for 30 minutes, using a desiccator. The mixture was poured on the master (described above) and evacuated again for an hour. The PDMS was subsequently cured in the oven, at 60°C, for 1-2 hours. After letting it harden overnight, the chips were cut in individual pieces using a razor. Holes were punched in the appropriate places using a 0.75 mm diameter biopsy punch (World Precision Instruments). The chip was cleaned using sticky tape and subsequently immobilized on a glass slide, by using plasma cleaning (Harrick Plasma, 29.6W, 60 s). Finally, the sealed PDMS microfluidic worm chips were cured at 80°C for two hours.

### Chemosensory stimuli

The microfluidic experiments were performed using M13 buffer. The following chemosensory stimuli were used, all diluted in M13: 0.1% sodium dodecyl sulfate (SDS), 10 mM copper sulfate (CuSO_4_), 1M or 0.5M glycerol (see below) and 0.25M sodium chloride (NaCl). The used concentrations of these stimuli were previously reported to act as repellents.^14^ This was confirmed by a behavioral assay (Movie S2), as described previously on the amphid^7^ or phasmid neurons^8^, before use in the microfluidics chip.

For stimulations with glycerol we used 1M for the calcium imaging experiments and 0.5M for the IFT imaging experiments, because the latter concentration evoked the same response as the first, but slightly less severe, making it easier to qualitatively describe the effect. Calcium imaging experiments with 0.5M glycerol resulted in similar results as 1M glycerol (Figure S3A).

### Microfluidic experiments

An adult worm was anesthetized in 5 mM levamisole for 10 min. Next, the worm was sucked up by a stainless steel pin (New England small tube corporation, 0.025’’ OD x 0.013’’ ID) into a Tygon tube connected to a 3 mL syringe filled with M13 buffer. The steel pin was inserted in the inlet of the worm channel and the worm was injected in the worm trap, tail-first, by manually applying pressure to the syringe.

The four fluidic inlets of the chip were connected with stainless steel pins and Tygon tubing to four 3 mL syringes filled with the appropriate solutions. The pressures in the syringes were controlled by a flow control system (Fluigent MFCS(tm)-EZ) and included software (Fluigent All-in-one 2019). By using a ‘four-flow system’, the flow directions of stimulus (channel 2) and buffer (channel 3) were controlled by the flows of fluorescein in channel 1 and 4, as described by Chronis et al. (2007).^17^ Prior to every experiment the pressures were calibrated to obtain appropriate flows.

### Imaging

Microscopic images were acquired using a custom-built epi-illuminated wide-field fluorescence set-up.^24,43^ All calcium imaging experiments were performed using a 40x air objective (Nikon, Plan Apo air 40x, N.A.: 0.95). Because the PHA and PHB neurons do not show GCaMP6s fluorescence when the worm is exposed to buffer, we had no prior knowledge about the location of these soma. We therefore made Z-stacks, in most experiments consisting of 5-10 frames, depending on the orientation of the worm, with a step size of 1 μm, to visualize the GCaMP6s intensity in one or more of the phasmid chemosensory neurons when exposing the worm tail to a stimulus. In most cases we could see an increase in fluorescence intensity in the dendrites that connect the cilia and the soma, which correlated with the neuronal activity (Figure 1B).

Depending on the lateral orientation of the worm, we could in some experiments draw intensity time traces of 2 or 3 somas, while in other experiments only 1 soma was in focus and suitable for measuring the intensity time traces. We therefore indicated n=x-y in the figure legends, in which x is the number of somas and y the number of nematodes.

For imaging IFT reporter strains, we used a 100x oil immersion objective (Nikon, CFI Apo TIRF 100x, N.A.: 1.49) with an additional 1.5x magnification (inside the microscope body).

Two color imaging, for the worm strain expressing kap-1::egfp and osm-3::mcherry, was performed by alternatingly switching between the 491nm and 561nm laser lines to image the eGFP and mCherry signals, respectively. The two color channels were imaged side by side on the camera chip using an Cairn OptoSplit II emission image splitter.

Imaging the cilium when exposing the worm to a repellent, is difficult. In many experiments the worm moved when it was exposed to the repellent, although we sedated the worms prior to the experiment (see above). The slightest movement, either in the Z-direction or when the worm tried to escape the stimulus by moving back in the worm trap, brought the cilium out of focus or field of view. In the figure legends we indicated the number of animals for which we have observed a qualitatively similar response (including the ones that moved too much to make a good kymograph). Typically of one third of those it was possible to make a kymographof quantitatively and qualitatively similar high-quality (i.e. with limited animal motion) as the kymograph shown in the figures..

### Analysis calcium imaging

For the calcium imaging experiments, we selected one or more phasmid chemosensory neurons (see above) and measured the mean intensity over the selected region of interest per subsequent Z projection. We used the average time the microscope needed to complete the Z-stack as time between subsequent Z-projections. The time was normalized such that t=0 equals the onset of the stimulus. GCaMP6s fluorescence intensity (ΔF/F_0_) was normalized by setting the highest fluorescence intensity in every individual measurement (F_max_) to 1, using custom written Matlab (2019a) scripts.

To compare the GCaMP6s fluorescence intensity profiles in response to the four chemosensory stimuli, we synchronized the profiles in such a way that the initial increase in fluorescence intensity at the start of the stimulus was set to t=0. We calculated centroids for time intervals of 30 seconds. To compare the traces in response to the four different stimuli, we performed an ANOVA test with multiple comparisons for the position of the centroids on the time and intensity axes.

### Kymograph analysis

Kymographs were generated using the open source ImageJ plugin KymographClear.^29^

3D plots (fluorescence intensity plotted over space and time) were generated from kymographs by averaging the intensity profiles over 1 s (glycerol) and 7 s (SDS). Intensities were smoothed on 5 pixels (250 nm) along the long axis of the cilia. Intensity profiles were corrected by the fluctuations of the background outside the cilia. In case two phasmid cilia overlapped, intensity profiles were divided by 2-(1/(exp(x-4.5)/200)+1).^24^ From the 3D-plots (Figure S5A and S8A), we generated snapshots over time (Figure 2C and 3B).

### Data and Software availability

Raw data from Figures 1-5 were deposited on Mendeley at http://dx.doi.org/10.17632/55sfvxvcjm.1

## References

1. White JG, Southgate E, Thomson JN, Brenner S. The structure of the nervous system of the nematode Caenorhabditis elegans. Philos Trans R Soc Lond B Biol Sci. 1986;314(1165):1–340.

2. Cook SJ, Jarrell TA, Brittin CA, Wang Y, Bloniarz AE, Yakovlev MA, et al. Whole-animal connectomes of both Caenorhabditis elegans sexes. Nature. 2019;571(7763):63–71.

3. Bargmann CI. Chemosensation in C. elegans. WormBook : the online review of C elegans biology. 2006:1–29.

4. Ben Arous J, Tanizawa Y, Rabinowitch I, Chatenay D, Schafer WR. Automated imaging of neuronal activity in freely behaving Caenorhabditis elegans. Journal of neuroscience methods. 2010;187(2):229–34.

5. Clark DA, Gabel CV, Gabel H, Samuel AD. Temporal activity patterns in thermosensory neurons of freely moving Caenorhabditis elegans encode spatial thermal gradients. The Journal of neuroscience : the official journal of the Society for Neuroscience. 2007;27(23):6083–90.

6. Faumont S, Rondeau G, Thiele TR, Lawton KJ, McCormick KE, Sottile M, et al. An image-free opto-mechanical system for creating virtual environments and imaging neuronal activity in freely moving Caenorhabditis elegans. PloS one. 2011;6(9):e24666.

7. Hilliard MA, Bargmann CI, Bazzicalupo P. C. elegans responds to chemical repellents by integrating sensory inputs from the head and the tail. Curr Biol. 2002;12(9):730–4.

8. Park J, Knezevich PL, Wung W, O’Hanlon SN, Goyal A, Benedetti KL, et al. A conserved juxtacrine signal regulates synaptic partner recognition in Caenorhabditis elegans. Neural development. 2011;6:28.

9. Ferkey DM, Sengupta P, L’Etoile ND. Chemosensory signal transduction in Caenorhabditis elegans. Genetics. 2021;217(3).

10. Goodman MB, Hall DH, Avery L, Lockery SR. Active currents regulate sensitivity and dynamic range in C. elegans neurons. Neuron. 1998;20(4):763–72.

11. Nguyen JP, Linder AN, Plummer GS, Shaevitz JW, Leifer AM. Automatically tracking neurons in a moving and deforming brain. PLoS Comput Biol. 2017;13(5):e1005517.

12. Nguyen JP, Shipley FB, Linder AN, Plummer GS, Liu M, Setru SU, et al. Whole-brain calcium imaging with cellular resolution in freely behaving Caenorhabditis elegans. Proc Natl Acad Sci U S A. 2016;113(8):E1074–81.

13. Dana H, Sun Y, Mohar B, Hulse BK, Kerlin AM, Hasseman JP, et al. High-performance calcium sensors for imaging activity in neuronal populations and microcompartments. Nat Methods. 2019;16(7):649–57.

14. Hilliard MA, Apicella AJ, Kerr R, Suzuki H, Bazzicalupo P, Schafer WR. In vivo imaging of C. elegans ASH neurons: cellular response and adaptation to chemical repellents. Embo j. 2005;24(1):63–72.

15. Kato S, Xu Y, Cho CE, Abbott LF, Bargmann CI. Temporal responses of C. elegans chemosensory neurons are preserved in behavioral dynamics. Neuron. 2014;81(3):616–28.

16. Zou W, Cheng H, Li S, Yue X, Xue Y, Chen S, et al. Polymodal Responses in C. elegans Phasmid Neurons Rely on Multiple Intracellular and Intercellular Signaling Pathways. Sci Rep. 2017;7:42295.

17. Chronis N, Zimmer M, Bargmann CI. Microfluidics for in vivo imaging of neuronal and behavioral activity in Caenorhabditis elegans. Nat Methods. 2007;4(9):727–31.

18. Singla V, Reiter JF. The primary cilium as the cell’s antenna: signaling at a sensory organelle. Science. 2006;313(5787):629–33.

19. Scholey JM. Intraflagellar transport. Annu Rev Cell Dev Biol. 2003;19:423–43.

20. Rosenbaum JL, Witman GB. Intraflagellar transport. Nature reviews Molecular cell biology. 2002;3(11):813–25.

21. Prevo B, Scholey JM, Peterman EJG. Intraflagellar transport: mechanisms of motor action, cooperation, and cargo delivery. The FEBS journal. 2017;284(18):2905–31.

22. Pan X, Ou G, Civelekoglu-Scholey G, Blacque OE, Endres NF, Tao L, et al. Mechanism of transport of IFT particles in C. elegans cilia by the concerted action of kinesin-II and OSM-3 motors. The Journal of cell biology. 2006;174(7):1035–45.

23. Oswald F, Prevo B, Acar S, Peterman EJG. Interplay between Ciliary Ultrastructure and IFT-Train Dynamics Revealed by Single-Molecule Super-resolution Imaging. Cell Rep. 2018;25(1):224–35.

24. Prevo B, Mangeol P, Oswald F, Scholey JM, Peterman EJ. Functional differentiation of cooperating kinesin-2 motors orchestrates cargo import and transport in C. elegans cilia. Nat Cell Biol. 2015;17(12):1536–45.

25. Mijalkovic J, van Krugten J, Oswald F, Acar S, Peterman EJG. Single-Molecule Turnarounds of Intraflagellar Transport at the C. elegans Ciliary Tip. Cell Rep. 2018;25(7):1701-7.e2.

26. Salzberg Y, Pechuk V, Gat A, Setty H, Sela S, Oren-Suissa M. Synaptic Protein Degradation Controls Sexually Dimorphic Circuits through Regulation of DCC/UNC-40. Curr Biol. 2020;30(21):4128-41.e5.

27. Mukhopadhyay S, Badgandi HB, Hwang SH, Somatilaka B, Shimada IS, Pal K. Trafficking to the primary cilium membrane. Molecular biology of the cell. 2017;28(2):233–9.

28. Schafer JC, Haycraft CJ, Thomas JH, Yoder BK, Swoboda P. XBX-1 encodes a dynein light intermediate chain required for retrograde intraflagellar transport and cilia assembly in Caenorhabditis elegans. Molecular biology of the cell. 2003;14(5):2057–70.

29. Mangeol P, Prevo B, Peterman EJ. KymographClear and KymographDirect: two tools for the automated quantitative analysis of molecular and cellular dynamics using kymographs. Molecular biology of the cell. 2016;27(12):1948–57.

30. Snow JJ, Ou G, Gunnarson AL, Walker MR, Zhou HM, Brust-Mascher I, et al. Two anterograde intraflagellar transport motors cooperate to build sensory cilia on C. elegans neurons. Nat Cell Biol. 2004;6(11):1109–13.

31. Qin H, Burnette DT, Bae YK, Forscher P, Barr MM, Rosenbaum JL. Intraflagellar transport is required for the vectorial movement of TRPV channels in the ciliary membrane. Curr Biol. 2005;15(18):1695–9.

32. van Krugten J, Danné N, Peterman EJG. TRPV channel OCR-2 is distributed along <em>C. elegans</em> chemosensory cilia by diffusion in a local interplay with intraflagellar transport. bioRxiv. 2020:2020.11.19.390005.

33. Mijalkovic J, Girard J, van Krugten J, van Loo J, Zhang Z, Loseva E, et al. Cutting off ciliary protein import: intraflagellar transport after dendritic femtosecond-laser ablation. Molecular biology of the cell. 2020;31(5):324–34.

34. Ezcurra M, Tanizawa Y, Swoboda P, Schafer WR. Food sensitizes C. elegans avoidance behaviours through acute dopamine signalling. Embo j. 2011;30(6):1110–22.

35. Pereira L, Kratsios P, Serrano-Saiz E, Sheftel H, Mayo AE, Hall DH, et al. A cellular and regulatory map of the cholinergic nervous system of C. elegans. Elife. 2015;4.

36. Denis V, Cyert MS. Internal Ca(2+) release in yeast is triggered by hypertonic shock and mediated by a TRP channel homologue. The Journal of cell biology. 2002;156(1):29–34.

37. Yu J, Yang W, Liu H, Hao Y, Zhang Y. An Aversive Response to Osmotic Upshift in Caenorhabditis elegans. eNeuro. 2017;4(2).

38. Mijalkovic J, van Krugten J, Oswald F, Acar S, Peterman EJG. Single-Molecule Turnarounds of Intraflagellar Transport at the C. elegans Ciliary Tip. Cell Rep. 2018;25(7):1701-7.e2.

39. Burghoorn J, Dekkers MP, Rademakers S, de Jong T, Willemsen R, Jansen G. Mutation of the MAP kinase DYF-5 affects docking and undocking of kinesin-2 motors and reduces their speed in the cilia of Caenorhabditis elegans. Proc Natl Acad Sci U S A. 2007;104(17):7157–62.

40. Woldemariam S, Nagpal J, Hill T, Li J, Schneider MW, Shankar R, et al. Using a Robust and Sensitive GFP-Based cGMP Sensor for Real-Time Imaging in Intact Caenorhabditis elegans. Genetics. 2019;213(1):59–77.

41. Frøkjaer-Jensen C, Davis MW, Hopkins CE, Newman BJ, Thummel JM, Olesen SP, et al. Single-copy insertion of transgenes in Caenorhabditis elegans. Nature genetics. 2008;40(11):1375–83.

42. Mijalkovic J, Prevo B, Oswald F, Mangeol P, Peterman EJ. Ensemble and single-molecule dynamics of IFT dynein in Caenorhabditis elegans cilia. Nature communications. 2017;8:14591.

43. van Krugten J, Peterman EJG. Single-Molecule Fluorescence Microscopy in Living Caenorhabditis elegans. Methods in molecular biology (Clifton, NJ). 2018;1665:145–54.

