## Supplementary information for "Differentiated dynamic response in *C. elegans* chemosensory cilia"

**Movie S1: Related to Figure 1. Movies used to quantify GCaMP6s fluorescence intensity over time.**

Representative movie of the maximum projection of a Z-stack of a worm tail expressing pan-neuronal GCaMP6s during stimulus exposure. Somas of PHA/PHB neurons light up at the start of the stimulus.

A) Exposure to 10mM $CuSO_4$. On- and offset of stimulus are indicated in the movie. Duration: 30 seconds. (Same data as in Figure 1G)

B) Exposure to 0.25M NaCl. On- and offset of stimulus are indicated in the movie. Duration: 30 seconds. (Same data as in Figure 1E)

**Movie S2: Related to Figure 1. *C. elegans* performs avoidance behavior in response to repellents.**

Movies of drop tests on young adult *C. elegans* with 0.1% SDS. A) Drop test was performed on young adult worms crawling forward on an NGM plate. A droplet of a repellent solution was placed in front of the worm, by gently touching the agar with a glass needle filled with the repellent solution, without touching the worm. B) Young adult worms crawling forward on an NGM plate were gently touched with an eyelash to induce backward motion. A droplet of repellent solution was placed near the tail, while the worm crawls backwards.

Crawling behavior of the animal was monitored after the worm came in contact with the droplet and recorded on Nikon SMZ1000.

### Figure S3

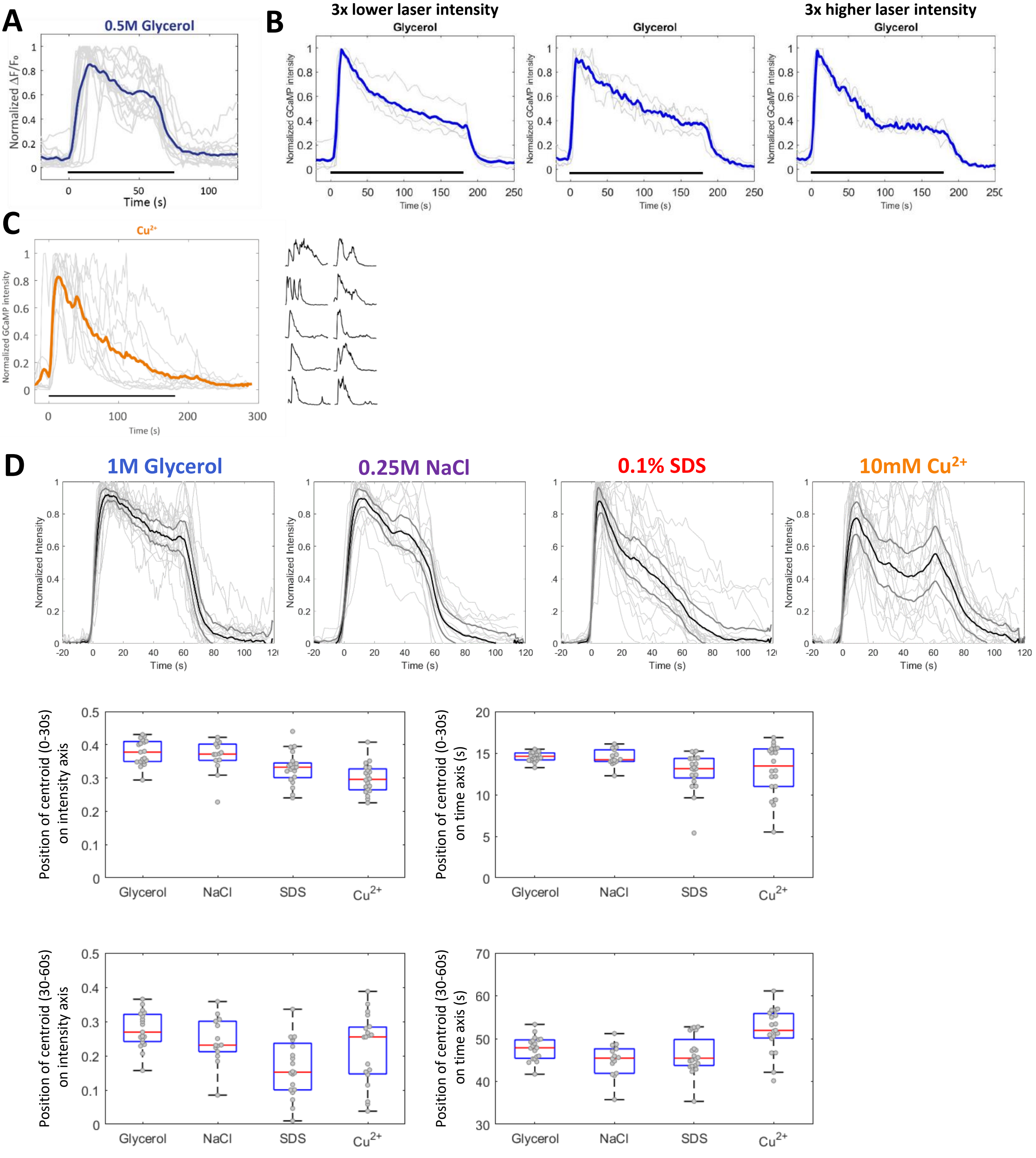


| P-values of centroids | Intensity 0-30s | Time 0-30s | Intensity 30-60s | Time 30-60s |
|---|---|---|---|---|
| Glycerol vs NaCl | 0.99 | 0.98 | 0.98 | 0.46 |
| Glycerol vs SDS | 0.02* | 0.03* | 0.00*** | 0.88 |
| Glycerol vs $Cu^{2+}$ | 0.00*** | 0.13 | 0.24 | 0.01** |
| NaCl vs SDS | 0.02* | 0.03* | 0.01** | 0.86 |
| NaCl vs $Cu^{2+}$ | 0.00*** | 0.10 | 0.56 | 0.00*** |
| SDS vs $Cu^{2+}$ | 0.19 | 0.93 | 0.11 | 0.00*** |

**Figure S3: Related to Figure 1.**

**A) GCaMP6s fluorescence intensity profiles in response to 0.5M glycerol.**

Normalized GCaMP6s intensity profiles in the PHA/PHB somas in response to 0.5M glycerol in M13. Light grey lines represent individual measurements, the thick blue line is the average. Black horizontal bar indicates the onset and duration (60 seconds) of the stimulus. n=16-11; n=x-y with x the number of somas and y the number of animals (see Methods).

**B) GCaMP6s fluorescence intensity profiles in response to long glycerol stimulus.**

Normalized GCaMP6s intensity profiles in the PHA/PHB somas in response to 1M glycerol in M13. Light grey lines represent individual measurements, the thick blue line is the average. Black horizontal bars indicate onset and duration (3 minutes) of the stimulus. Middle panel: measurements made with the imaging settings that are used in the rest of this study, n=4-3; left panel represents measurements made with 3x lower laser intensity, n=4-3; right panel with 3x higher laser intensity, n=3-2.

**C) GCaMP6s intensity profiles in response to long $Cu^{2+}$ stimulus.**

Normalized GCaMP6s intensity profiles in the PHA/PHB somas in response to 10 mM $Cu^{2+}$ in M13. Light grey lines represent individual measurements, thick orange line the average. Black horizontal bar indicates onset and duration (3 minutes) of the stimulus. n=10-5. On the right the individual intensity profiles are depicted separately.

**D) Statistical analysis of GCaMP6s intensity profiles in response to 60 seconds exposure to 1M glycerol; 0.25M NaCl; 0.1%SDS and 10mM $Cu^{2+}$.**

Upper graphs show normalized GCaMP6s intensity profiles in the PHA/PHB somas in response to the four different chemosensory stimuli, with synchronized time. Black lines indicate the mean, grey thick lines represent the standard deviation, grey thin lines represent the individual measurements. (Same measurements as in Figure 1C-F.)

Boxplots show position of the centroids of the GCaMP6s intensity profile, on the intensity axis (boxplots on the left) and time axis (boxplots on the right) in response to the stimulus depicted on the x-axis. Upper boxplots represent first time interval (0-30s); lower boxplots the second time interval (30-60s).

The table below shows the P-values of the centroids depicted in the boxplots, determined by ANOVA with multiple comparisons. The P-values shown here are used to calculate the number of asterisks shown in Figure 1G. ns, not significant; * $p<0.05$; ** $p<0.01$; *** $p<0.001$.

**Movie S4: Related to Figure 2. Response of IFT machinery to glycerol stimulus.**

Representative movies of worm strains expressing fluorescently labeled IFT-components during glycerol exposure (30 seconds).

A) KAP-1::eGFP (odd frames; left side of the camera); OSM-3::mCherry (even frames; right side of the camera). Exposure time 75 ms. t=00:08 start glycerol stimulus; t=00:38; end of glycerol stimulus.

B) XBX-1::eGFP. Exposure time 150 ms. t=00:20 start glycerol stimulus; t=00:50 end of glycerol stimulus.

C) OSM-6::eGFP. Exposure time 150 ms. t=00:20 start glycerol stimulus; t=00:50 end of glycerol stimulus.

### Figure S5

**A**

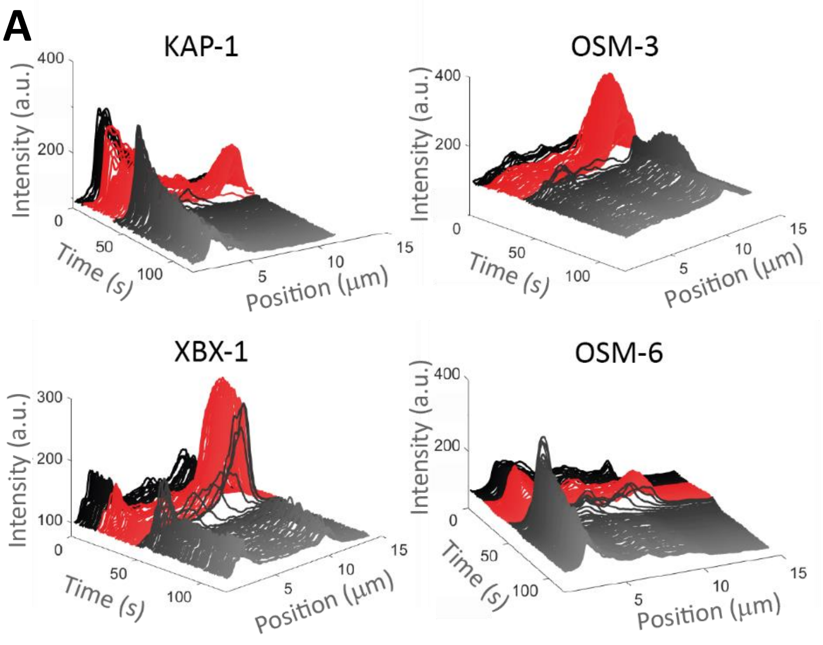


**B**

**0.5M Glycerol**

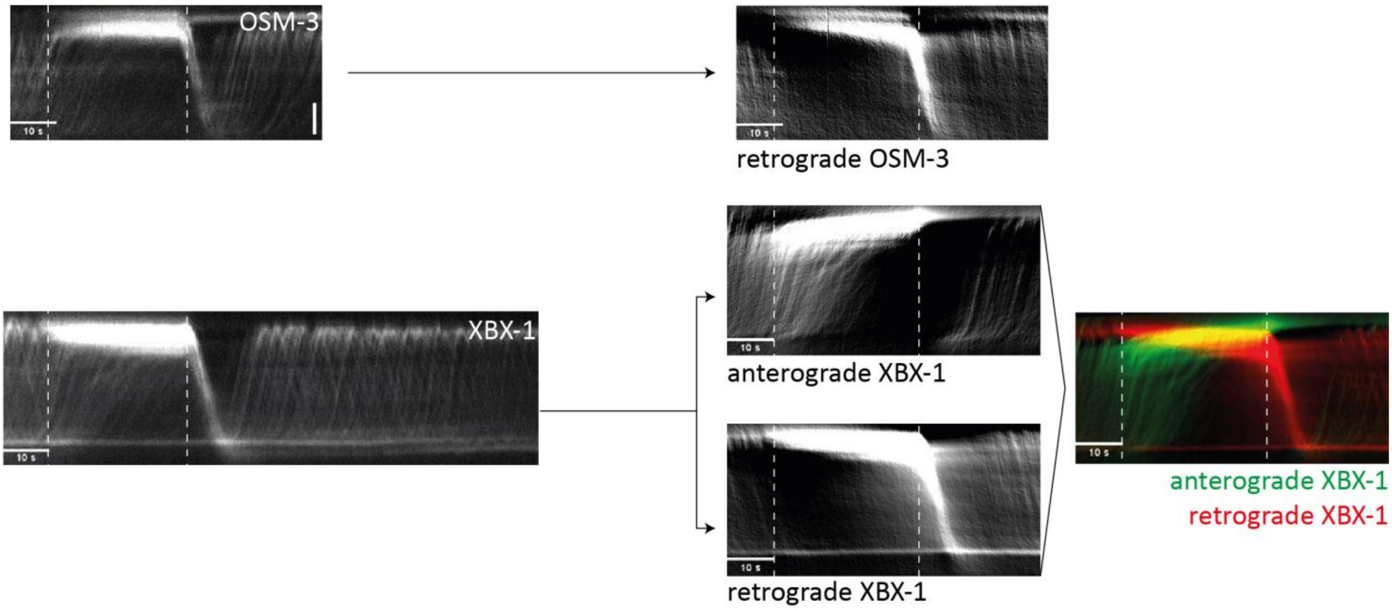


**Figure S5: Related to Figure 2.**

**A) 3D plots of the IFT machinery in response to glycerol.**

3D plots (fluorescence intensity plotted over space and time) of the IFT machinery before, during and after 0.5M glycerol exposure. 3D plots were generated from kymographs (depicted in Figure 2D) by averaging the intensity profiles over 1 s. Intensities were smoothed on 5 pixels (250 nm) along the long axis of the cilia. Black lines indicate the situation before the stimulus; red lines during the stimulus and grey lines after.

**B) Fourier-filtered kymographs of OSM-3 and XBX-1 during glycerol exposure.**

Retrograde Fourier-filtered kymograph of OSM-3::mCherry, and anterograde and retrograde Fourier-filtered kymograph of XBX-1::eGFP in response to 0.5M glycerol (non-filtered kymographs on the left are shown in Figure 2D). Right kymograph shows the overlay of anterograde (green) and retrograde (red) moving XBX-1. Vertical: position in cilium, scale bar: 2 μm; horizontal: time. First dotted line indicates the start of the stimulus, second dotted line the end of the stimulus.

### Figure S6

**0.5M Glycerol**

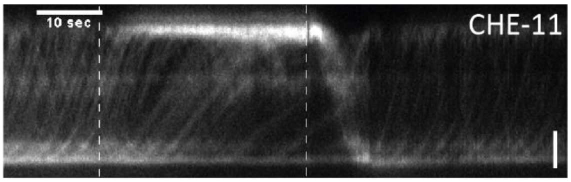


**0.1% SDS**

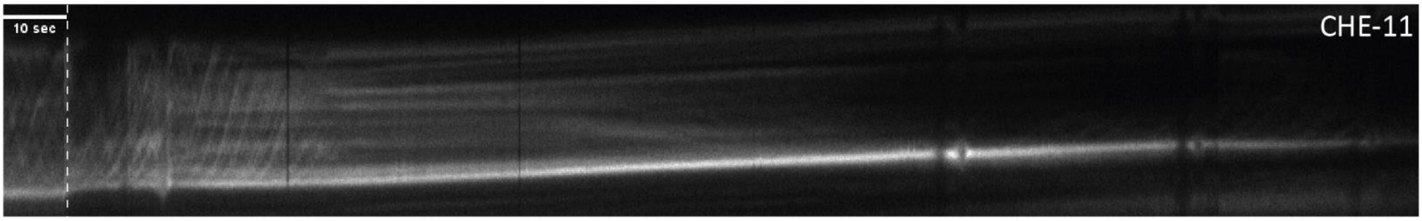


**Figure S6: Related to Figure 2 and 3. IFT-A in response to glycerol and SDS.**
Kymographs of CHE-11::mCherry (IFT-A) in response to 0.5M glycerol (upper kymograph), n=7; and 0.1% SDS (lower kymograph), n=4. Vertical: position in cilium, scale bar: 2 µm; horizontal: time. First dotted line indicates the start of the stimulus, second dotted line (in case of glycerol) end of stimulus. Representative kymographs of n=x animals for which we have observed a qualitatively similar response (see Methods).

### Figure S7

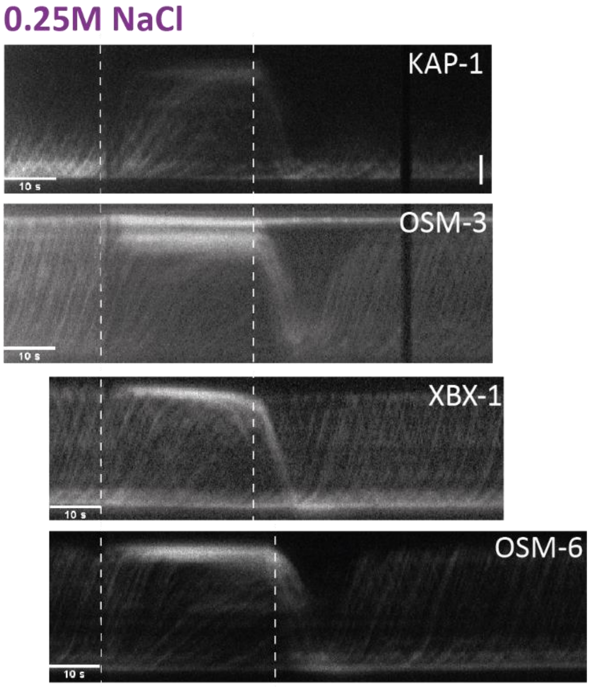


**Figure S7: Related to Figure 2. Response of IFT machinery to exposure to high concentrations of salt.**

Kymographs of KAP-1::eGFP (kinesin-II), n=3; OSM-3::mCherry, n=3; XBX-1::eGFP (IFT dynein), n=3; and OSM-6::eGFP (IFT-B), n=4 upon 0.25M NaCl exposure. Vertical: position in cilium, scale bar: 2 µm; horizontal: time. First dotted line indicates the start of the stimulus, second dotted line end of stimulus. Representative kymographs of n=x animals for which we have observed a qualitatively similar response (see Methods).

### Figure S8

**A**

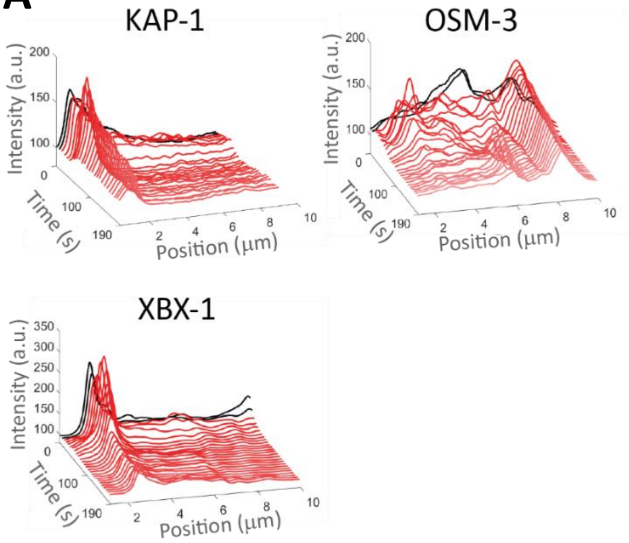


**B** 10mM $Cu^{2+}$

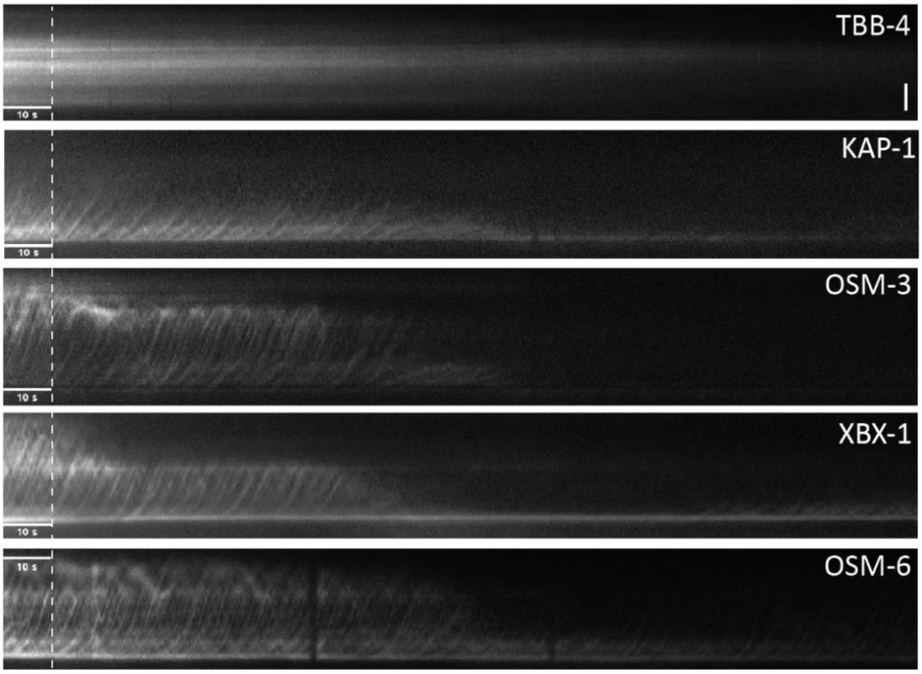


**Figure S8: Related to Figure 3.**

**A) 3D plots of the IFT machinery in response to SDS.**

3D plots (fluorescence intensity plotted over space and time) of the IFT machinery before and during 0.1% SDS exposure. 3D plots were generated from kymographs (shown in Figure 3C) by averaging the intensity profiles over 7 s. Intensities were smoothed on 5 pixels (250 nm) along the long axis of the cilia. Black curves indicate the situation before the stimulus; red curves during the stimulus.

**B) Response of IFT machinery to 10 mM $Cu^{2+}$ exposure.**

Kymographs of TBB-4::eGFP (tubulin), n=8; KAP-1::eGFP (kinesin-II), n=6; OSM-3::mCherry, n=6; XBX-1::eGFP (IFT dynein), n=10; and OSM-6::eGFP (IFT-B), n=8; upon 10 mM $Cu^{2+}$ exposure. Vertical: position in cilium, scale bar: 2 µm; horizontal: time. Dotted line indicates the start of the stimulus. Representative kymographs of n=x animals for which we have observed a qualitatively similar response (see Methods).
